# Odd skipped-related 1 controls the pro-regenerative response of Fibro-Adipogenic Progenitors

**DOI:** 10.1101/2022.07.04.498663

**Authors:** Georgios Kotsaris, Taimoor H. Qazi, Christian H. Bucher, Sophie Pöhle-Kronawitter, Vladimir Ugorets, William Jarassier, Stefan Börno, Bernd Timmermann, Claudia Giesecke-Thiel, Pedro Vallecillo-García, Aris N. Economides, Fabien Le Grand, Petra Knaus, Sven Geissler, Sigmar Stricker

## Abstract

Skeletal muscle regeneration requires the coordinated interplay of diverse tissue-resident- and infiltrating cells. Fibro-adipogenic progenitors (FAPs) are an interstitial cell population that provides a beneficial microenvironment for muscle stem cells (MuSCs) during muscle regeneration. Here we show that the transcription factor Osr1 is essential for FAPs to communicate with MuSCs and infiltrating macrophages, thus coordinating muscle regeneration. Conditional inactivation of Osr1 impaired muscle regeneration with reduced myofiber growth and formation of excessive fibrotic tissue with reduced stiffness. Osr1-deficient FAPs acquired a fibrogenic identity with altered matrix secretion and cytokine expression resulting in impaired MuSC viability, expansion and differentiation. Immune cell profiling suggested a novel role for Osr1-FAPs in macrophage polarization. *In vitro* analysis suggested that increased TGFβ signaling and altered matrix deposition by Osr1-deficient FAPs actively suppressed regenerative myogenesis. In conclusion, we show that Osr1 is central to FAP function orchestrating key regenerative events such as inflammation, matrix secretion and myogenesis.

## Introduction

Skeletal muscle, which accounts for 30-40% of the total body mass in mammals, is one of the few tissues capable of scarless healing. However, large-scale trauma or muscle disease often lead to replacement of damaged tissue by fibrous and fatty infiltrates ^1^. The regenerative capacity of skeletal muscle depends on tissue resident muscle stem cells (MuSCs) that are essential for skeletal muscle regeneration ^2,3^. MuSCs are rapidly activated, and upon tissue damage start to proliferate and give rise to a progenitor pool capable of replacing damaged muscle fibers ^4,5^. During regeneration, myogenesis is tightly connected to an interplay of multiple other cell types. Skeletal muscle regeneration follows the general principle of wound healing and begins with an initial inflammatory response characterized by infiltration of immune cells, which represent the first wave of cells expanding in the injured area ^6^. Among immune cells, macrophages play a key role in orchestrating the regeneration process. Pro-inflammatory (historically termed M1) macrophages are required for the clearance of tissue debris, attraction and modulation of further immune cells and activation of MuSCs ^7^. Subsequently these macrophages convert into anti-inflammatory (M2) phenotypes, which promotes muscle progenitor differentiation into myocytes and their fusion to new myofibers ^4,5,8,9^. Immune cell infiltration in the injury area is also associated with the immediate activation and expansion of tissue-resident stromal cells called fibro-adipogenic progenitors (FAPs) ^10^. FAPs, originally identified by expression of the cell surface markers Sca-1 or PDGFRα ^11,12^, generate a beneficial microenvironment for regeneration in part via secreted signaling molecules and via the formation of a transient extracellular matrix (ECM) ^11,13–15^. While FAPs are essential for effective muscle regeneration under physiological conditions ^3,16^, they are the source of fibrosis and fatty infiltration under degenerating or chronic inflammatory states ^14,17–21^. To control transitional FAP pool expansion and prevent fibro-fatty infiltration, pro-inflammatory macrophages induce FAP apoptosis via TNFα in mid-regeneration thus limiting transient ECM production and making way for regenerating muscle fibers ^22^. Conversely, anti-inflammatory M2 macrophages create a TGFβ-rich environment that induces differentiation of FAPs into myofibroblasts, which increasingly synthetize new ECM components ^15,22^. Accordingly, many myopathies, including amyotrophic lateral sclerosis and Duchenne muscular dystrophy, are associated with exacerbated inflammatory responses, resulting in deregulated FAP function, excessive fibrotic tissue formation, and loss of muscle function ^21,23^. Despite increasing knowledge about the role of FAPs during the regenerative process, no intrinsic transcriptional regulator is known that controls key aspects of their different functions. We previously identified the zinc finger transcription factor Odd skipped-related 1 (Osr1) as a key regulator of the pro-myogenic function in an embryonic FAP-like cell population, which also is a developmental source of adult FAPs ^24^. While *Osr1* reporter gene and protein expression was undetectable in homeostatic muscle, it was reactivated during muscle regeneration ^25^. Here we show that *Osr1* is required for FAP function during skeletal muscle regeneration. Loss of *Osr1* leads to a pro-fibrotic orientation of FAPs and impairs both FAP-macrophage and FAPs-MuSC communication networks resulting in impaired regenerative myogenesis and persistent fibrosis. This demonstrates that Osr1 is a key transcriptional regulator of FAP regenerative function protecting FAPs from assuming a detrimental pro-fibrotic and anti-myogenic state.

## Results

### Conditional inactivation of Osr1 impairs muscle regeneration

First, *Osr1* expression was analyzed in the single cell dataset from Oprescu et al. ^26^, confirming *Osr1* expression in FAPs throughout muscle regeneration (Fig. S1a) in agreement with our previous data ^25^. Osr1-flox mice generated from an *Osr1* multifunctional allele ^25^ were mated to CAGG-CreER animals to allow timed inactivation of *Osr1* based on tamoxifen delivery concomitant with injury (Fig. 1a). Cre-mediated recombination inverts a GFP cassette to be driven by the endogenous *Osr1* promoter (Fig. 1a). To induce recombination, tamoxifen was administered concomitant to freeze-pierce injury of the tibialis anterior muscle (TA), and on the following two days (Fig. 1b). Expression of GFP from the recombined *Osr1* allele was confirmed via flow cytometry exclusively in Lin- (non-hematopoietic, non-endothelial) cells (Fig. S1b). Osr1-deficient Osr1^flox/flox^;CAGG^CreER^ animals are further termed Osr1cKO. *Osr1* mRNA expression was efficiently decreased by 90% in Osr1cKO total muscle samples on 3 days post injury (dpi) (Fig. S1c).

**Figure 1.**
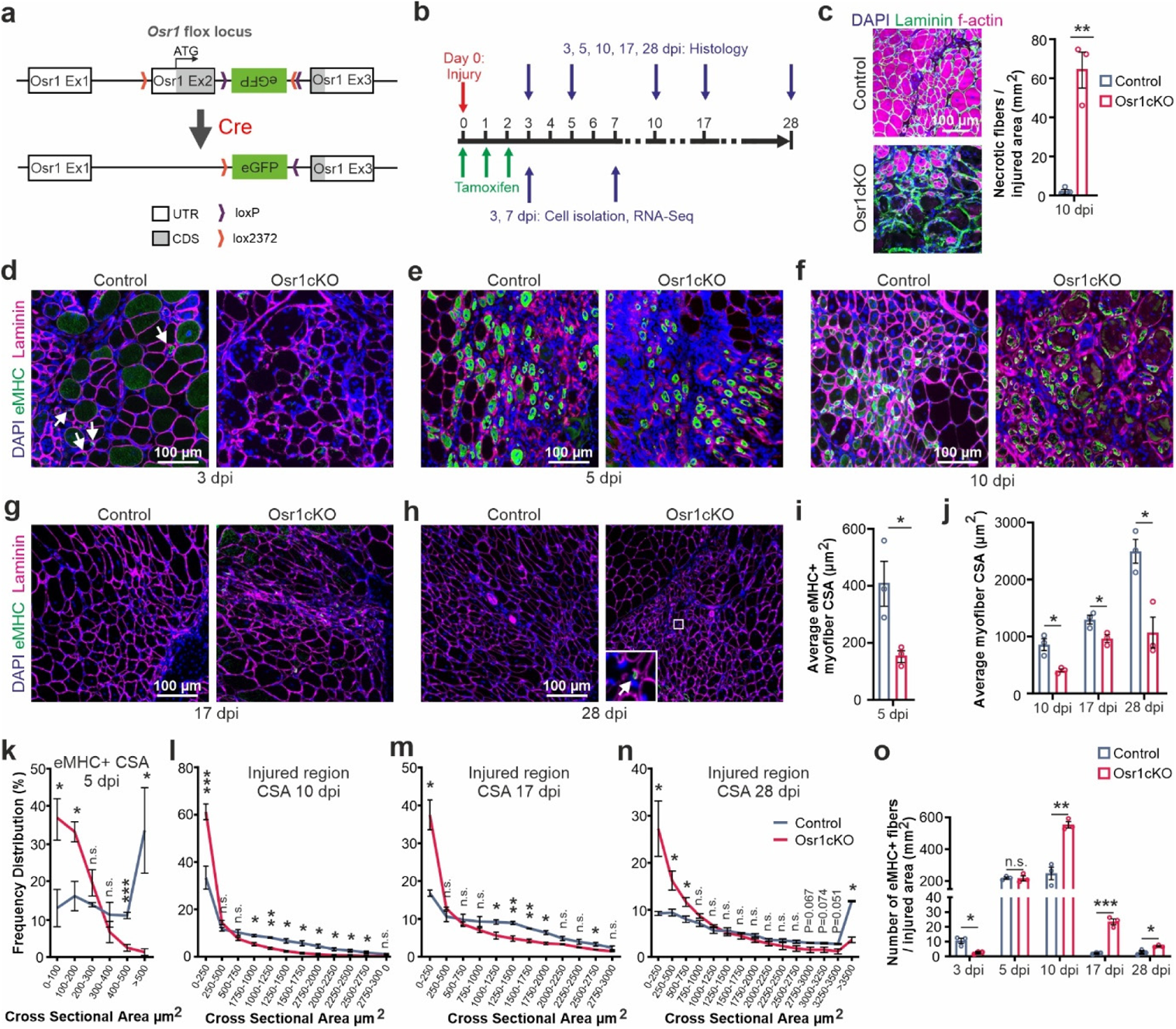
Conditional inactivation of Osr1 impairs muscle regeneration. **a** Schematic representation of the *Osr1* conditional allele and Cre-mediated locus recombination **b** Schematic depiction of Tamoxifen administration and analysis time points (dpi: days post injury). **c** Phalloidin labeling for f-actin at 10 dpi to detect necrotic fibers; quantification shown right (n=3). **d-h** Immunolabeling for embryonal myosin heavy chain (eMHC) and Laminin on control and Osr1cKO muscle sections at indicated dpi; nuclei were stained for DAPI. **I** Quantification of eMHC+ myofiber cross sectional area (CSA) in the injury region of control and Osr1cKO muscle at 5 dpi (n=3). **j** Quantification of myofiber cross sectional area (CSA) in the injury region of control and Osr1cKO muscle at 10, 17 and 28 dpi (n=3). **k** eMHC+ Myofiber size frequency distribution in the injury region of control and Osr1cKO muscle at 5 dpi (n=3). **l-n** Myofiber size frequency distribution in the injury region of control and Osr1cKO muscle at 10, 17 and 28 dpi (n=3). **o** Quantification of eMHC positive fibers per area at indicated dpi (n=3). Data are mean ± SEM; P-value calculated by two-sided unpaired t-test; * p < 0.05, **p < 0.01, ***p < 0.001. N-numbers indicate biological replicates (mice per genotype).

No major histological differences were observed between Osr1cKO and control muscles at 3 and 5 dpi (Fig. S2a). Fiber density and size decreased in injured regions of Osr1cKO as well as controls between 3 and 5 dpi (Fig. S2a), consistent with degeneration of injured muscle tissue driven by the initial pro-inflammatory phase. The injury area of Osr1cKO muscle showed pronounced accumulation of granulation tissue and abundance of necrotic fibers at 10 dpi compared to controls (Fig. S2a). Phalloidin staining, detecting filamentous actin of intact fibers, confirmed persistence of necrotic fibers in Osr1cKO animals at 10 dpi (Fig. 1c) while tissue necrosis was completely resolved in control muscles, which displayed a well-organized fiber structure.

Overall, regenerating myofibers appeared smaller and more variable in size in Osr1cKO muscle (Fig. S2a). Immunolabeling for laminin to outline muscle fibers and for embryonal myosin heavy chain (eMHC) to label newly regenerating fibers confirmed impaired post-injury myofiber growth in Osr1cKO animals (Fig. 1d-h). The average size (cross sectional area, CSA) of eMHC+ myofibers at 5 dpi was reduced in Osr1cKO muscle to less than 50% of control levels (Fig. 1i). At 10, 17 and 28 dpi myofibers in the injury area of Osr1cKO mice were 30-60% smaller compared to corresponding controls (Fig. 1j). While a steep increase in fiber size was observed in the controls especially between 17 and 28 dpi, fiber size remained constant in the Osr1cKO. Plotting discrete size windows (CSA distribution frequency) confirmed a shift towards extremely small fiber calibers in Osr1cKO muscle, with approx. 70% of eMHC+ fibers having a CSA of less than 200 µm^2^ at 5 dpi (Fig. 1k), and approx. 50% of fibers having a CSA less than 500 µm^2^ at 10, 17 and 28 dpi (Fig. 1l-n).

Quantification of actively regenerating eMHC+ fibers within the injury area showed a delay in formation and maturation of new fibers. Compared to the controls, Osr1cKO muscles exhibited a significantly lower amount of eMHC+ fibers at 3 dpi (Fig. 1d, o), which was equalized at 5 dpi (Fig. 1e, o). Conversely, a higher proportion of eMHC+ fibers was observed in Osr1cKO muscle at 10 dpi (Fig. 1f, o), which accumulated around apparently non-resolving necrotic fibers (Fig. 1f). At 17 and 28 dpi eMHC+ fibers had almost completely disappeared in controls but persisted in the Osr1cKO (Fig. 1g, h, o).

In conclusion, inactivation of Osr1 leads to impaired necrotic tissue resolution and delayed formation, maturation and growth of regenerating myofibers.

### Loss of Osr1 leads to reduced regenerating tissue stiffness and persistent fibrosis

Immunolabeling for laminin (Fig. 1c-g) and collagen type VI (Fig. S3a) indicated an increase in ECM in Osr1cKO muscle. Picrosirius red staining was performed to evaluate ECM deposition. No difference was observed at 3 dpi between control and Osr1cKO muscle, indicating the initial transient fibrotic response as a normal feature of the regeneration process (Fig. 2a, f). Starting at 5 dpi, a significant increase in ECM deposition was found in Osr1cKO muscle (Fig. 2b, f). While ECM remodeling in controls led to a marked decrease in picrosirius red staining between 5 and 10 dpi, it peaked at 10 dpi in Osr1cKO animals, and started to decrease between 10 and 17 dpi (Fig. 2c, d, f). At 17 and 28 dpi, Osr1cKO animals still showed approx. twice as much picrosirius red staining than controls (Fig. 2d, e, f) indicating persistent fibrosis. Fibrotic scarring is typically associated with increased tissue stiffness ^27^, as it is also observed in *mdx* animals used as a model for Duchenne muscular dystrophy ^28^. However, these observations regularly are made by assessing passive tissue stiffness at a late time point of disease ^29,30^. Nanoindentation was performed to assess the impact of altered tissue remodeling on the mechanical properties of the regenerating tissue during the initial phase of transient pro-regenerative ECM formation (Fig. 2g). Dynamic alterations in tissue stiffness were first defined in wild type mice by comparing the injured region of damaged muscles at 3, 4 and 5 dpi with the corresponding intact (contralateral) muscle. This showed a transient stiffness increase at 3 and 4 dpi, which decreased back to the range of uninjured levels already at 5 dpi (Fig. S3b). In regenerating Osr1cKO muscles no transient increase in stiffness was observed, but rather a tissue softening occurred at 3 dpi (Fig. 2h). Injured Osr1cKO muscle consistently exhibited lower stiffness than controls at 5 and even at 10 dpi (Fig. 2h). Taken together, loss of Osr1 resulted in a softened transient ECM during regeneration followed by persistent fibrosis.

**Figure 2.**
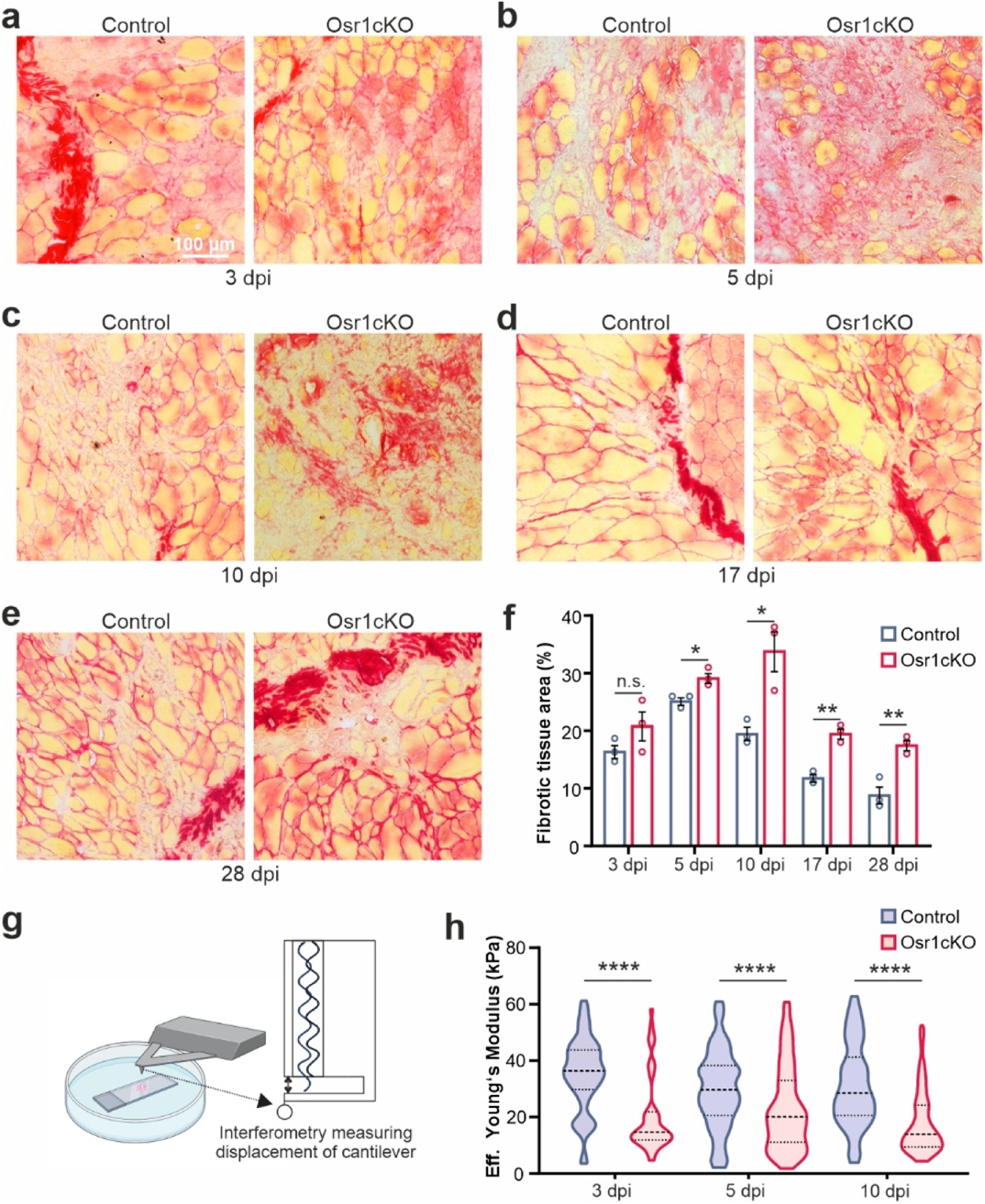
Loss of Osr1 results in persistent fibrosis and tissue softening. **a-e** Picrosirius Red staining on control and Osr1cKO muscle sections at indicated days post injury (dpi). **f** Quantification of picrosirius red staining (n=3). **g** Schematic representation of nanoindentation method to assess tissue stiffness. **h** Quantification of nanoindentation stiffness measurements of the injured area in muscle tissue sections of control and Osr1cKO mice (n=3). Violin plots show full data range, mean value, first and third quartiles are indicated. Data are mean ± SEM; *P*-value calculated by two-sided unpaired *t*-test; * p < 0.05, **p < 0.01, ****p < 0.0001. N-numbers indicate biological replicates (mice per genotype).

### Osr1-deficient FAPs have an intrinsic defect in pool expansion

We next assessed possible direct effects of Osr1 depletion on FAPs. Flow cytometric analysis showed decrease of FAP numbers in the whole Osc1cKO muscle by an approx. 7% compared to the control at 3 dpi (Fig. 3a). At 7 dpi, FAP numbers were not significantly altered (Fig. 3a). Considering the small lesion, this slight decrease in the whole muscle tissue suggests larger differences in the direct injury region. FAPs isolated by FACS showed approx. 80% decreased Ki67 labeling (Fig. 3b), and approx. three-fold increased apoptosis rate (Fig. 3c).

**Figure 3.**
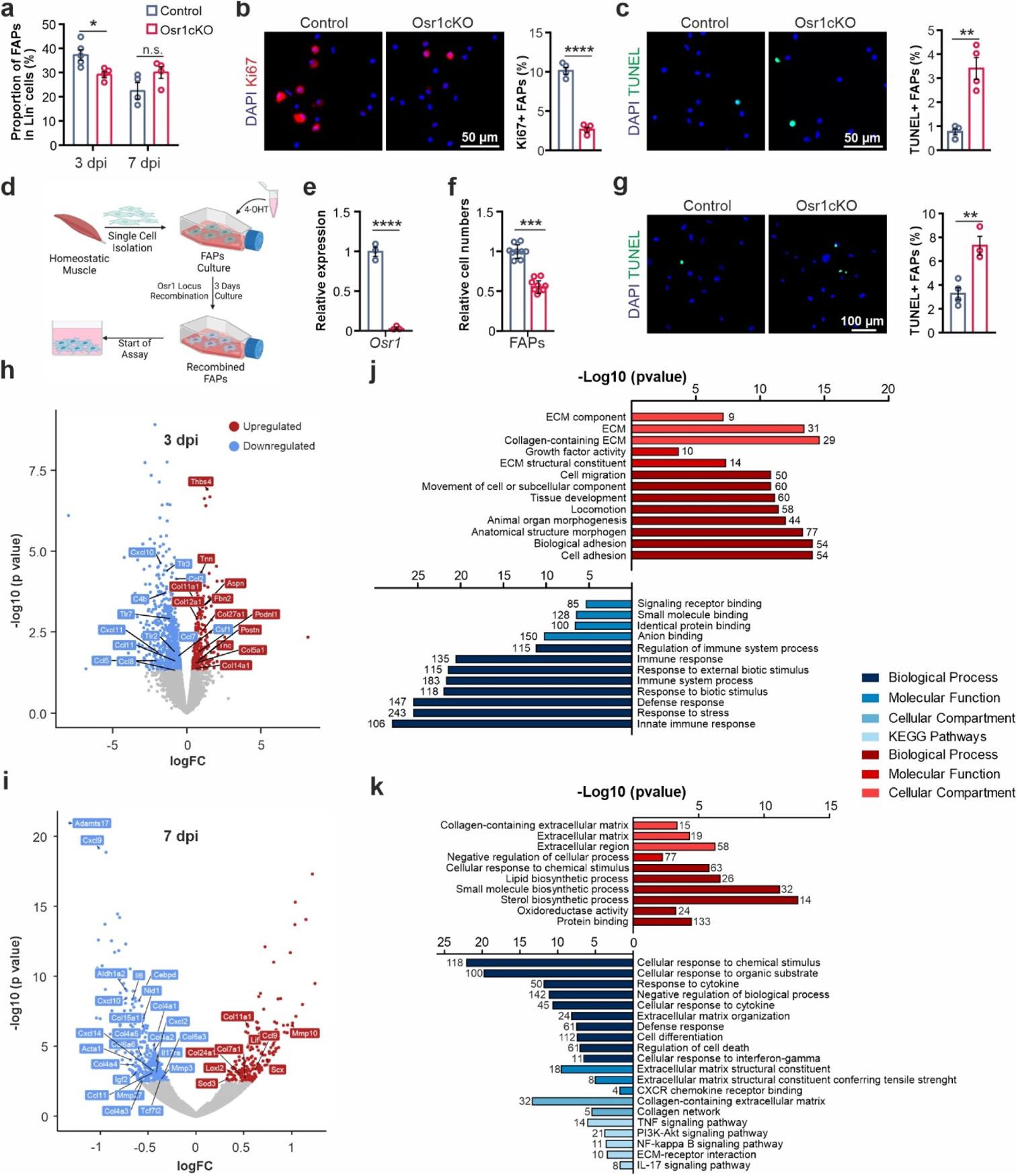
Osr1 deletion affects FAP expansion and transcriptomic response after acute injury. **a** Flow cytometry analysis of FAP numbers at 3 and at 7 dpi (at 3 dpi n=5 for control and n=4 for Osr1cKO, at 7 dpi n=4 each). **b** Ki67 labeling of 7 dpi FAPs isolated by FACS; quantification right (n=4). **c** TUNEL labeling 7 dpi FAPs isolated by FACS; quantification right (n=4). **d** Schematic depiction of FAP *in vitro* recombination. **e** RT-qPCR analysis of *Osr1* expression after *in vitro* recombination. **f** Cell numbers of *in vitro* recombined FAPs after 6 days of culture in growth medium. **g** TUNEL labeling on *in vitro* recombined FAPs after 4 days of culture in growth medium. **h, i** Volcano plot of DE genes between Osr1^flox/+^;CAGG-Cre^+^ (control) and Osr1cKO FAPs at 3 or 7 dpi. **j, k** GO analysis of genes upregulated (red) or downregulated (blue) in Osr1cKO FAPs at 3 or 7 dpi. Data are mean ± SEM; P-value calculated by two-sided unpaired t-test; * p < 0.05, **p < 0.01, ***p < 0.001, ****p < 0.0001. N-numbers indicate biological replicates (mice per genotype).

To assess whether these defects were cell autonomous, we performed *in vitro* recombination of the Osr1^flox/flox^ allele in FAPs isolated by pre-plating (Fig. 3d), yielding cells phenotypically and biochemically similar to FAPs isolated by FACS ^31^. 4-hydroxytamoxifen treatment resulted in an approx. 98% decrease of *Osr1* mRNA expression (Fig. 3e). In line with *in vivo* results, also these *in vitro* recombined FAPs showed significantly reduced cell numbers after 6 days of culture in growth medium (Fig. 3f) and increased apoptosis (Fig. 3g) suggesting that Osr1 is required for FAP viability.

### Osr1-deficient FAPs show altered cytokine and ECM gene expression profiles

To gain deeper insight into intrinsic effects of Osr1 depletion, we performed transcriptome analysis of FAPs isolated by FACS at 3 and 7 dpi. To enrich FAPs from the injury region, cells with active *Osr1* expression were isolated based on GFP expression of the recombined Osr1^flox^ allele. Efficient deletion of floxed exon 2 and consequent lack of exon 3 expression was confirmed in the RNA Sequencing (RNA-Seq) data (Fig. S4a). RNA-Seq revealed a total of 950 differentially expressed genes (DEG), of which 261 were upregulated and 689 were downregulated in homozygous Osr1cKO FAPs compared with heterozygous controls at 3 dpi (Fig. 3h). At 7 dpi, 544 DEGs were determined, of which 206 were up- and 338 were downregulated in Osr1cKO FAPs (Fig 3i). We then performed gene ontology (GO) annotation clustering of the DEGs according to cellular component, biological process and signaling pathway. This revealed that a significant number of DEGs upregulated in Osr1cKO FAPs at 3 dpi were associated with the ECM (GO:0031012), “Collagen-containing ECM” (GO: 0062023), “Extracellular structural constituent” (GO:0005201) or cell-matrix interaction such as “Cell adhesion” (GO:0007155) (Fig. 3j). Upregulated DEGs at 7 dpi were also mainly associated with terms related to the ECM (Fig. 3k). DEGs downregulated in Osr1cKO FAPs showed enrichment of genes related to the “innate immune response” (GO:0045087) or “regulation of immune system” (GO:0002682) at 3 dpi (Fig. 3j) and enrichment of genes belonging to “response to cytokine” (GO:0034097) and “defense response” (GO:0034097) at 7 dpi (Fig. 3k). Accordingly, analysis of genes commonly deregulated at both time points (total: 96 genes: 32 up- and 45 down-regulated at both time points, 19 genes with opposite regulation; Fig. S4b) confirmed that they were associated with GO terms related to “collagen-containing extracellular matrix” and “defense response,” respectively. (Fig. S4c, d).

In summary, loss of Osr1 impairs initial FAP expansion and leads to a sustained shift in their transcriptional profile. These transcriptional alterations might affect several key functions attributed to FAPs during regeneration ^6^, including the synthesis and remodeling of the transient ECM, but also their immunomodulatory properties.

### Loss of Osr1 disrupts FAP-immune cell interplay and prevents regenerative macrophage polarization

Since transcriptome analysis indicated altered immunomodulatory properties of Osr1cKO FAPs, we investigated possible implications for the interplay between FAPs and immune cells during muscle regeneration. Heatmap depiction confirmed a global downregulation of genes, including numerous secreted signalling molecules and cytokines, belonging to GO groups “Inflammatory response” and “Response to cytokine” in Osr1cKO FAPs (Fig. 4a, b).

**Figure 4.**
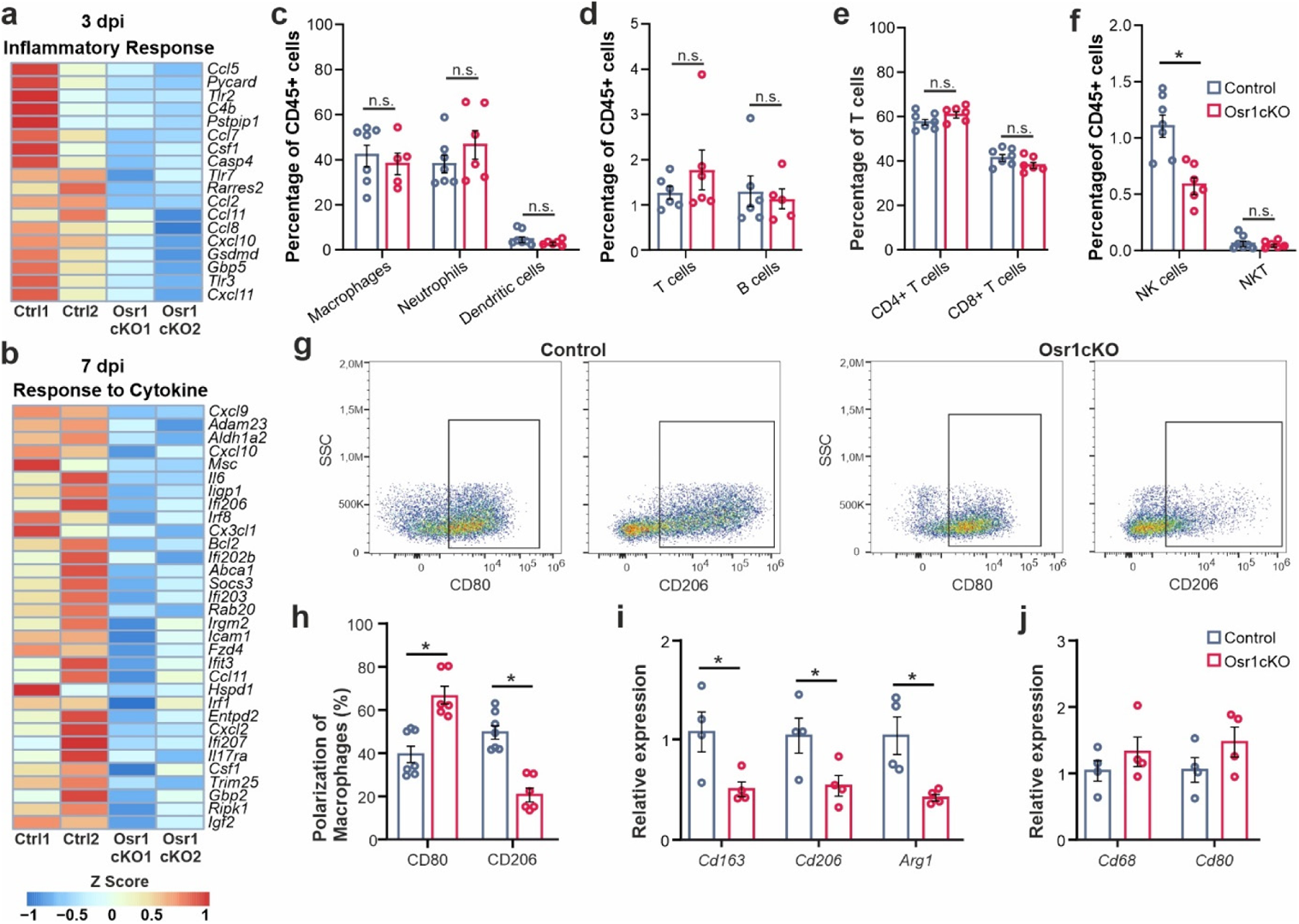
Osr1 is required for macrophage polarization during skeletal muscle regeneration. **a, b** Heat maps of DE genes in 3 or 7 dpi control vs. Osr1cKO FAPs belonging to GO terms “Inflammatory response” and “Response to Cytokine”. **c-f** Relative quantification of myeloid subpopulations by flow cytometry (c), overall T- and B-cells (d), CD4 and CD8 T cells (e), and NK/NKT cells (f) isolated from control or Osr1cKO 3 dpi muscle (n=7 for control and n=6 for Osr1cKO). **g** Representative FACS plots of CD80 / CD206 macrophage polarization analysis. **h** Quantification of CD80 and CD206 macrophages in control or Osr1cKO muscle at 3 dpi. **i, j** RT-qPCR analysis of *Cd163*, *Cd206* and *Arg1*, or *Cd68* and *Cd80* in whole muscle lysate (n=4). Data are mean ± SEM; P-value calculated by Mann-Whitney test in c-f and by two-sided unpaired t-test in h-j; *p < 0.05. N-numbers indicate biological replicates (mice per genotype).

Flow cytometry analysis of 3 dpi muscles revealed distinct local immune cell profiles in Osr1cKO and control animals, but confirmed the presence of major myeloid and lymphoid subpopulations in both specimen (Fig. S5a). Equal total cell numbers and CD45+ cell numbers were isolated from control or Osr1cKO muscle (Fig. S5b). No significant difference in the relative overall numbers of leukocytes, macrophages, neutrophils, dendritic cells (Fig. 4c, Fig. S5c), or dendritic cell subsets (Fig. S5d) was detected between Osr1cKO and control muscles. Numbers of B- and T-cells were globally unaltered (Fig. 4d), as were T-cell-subsets (Fig. 4e, S5e). This altogether argues against a global effect of Osr1 depletion in FAPs on immune cell infiltration. Low levels of circulating memory cells in the blood confirm that animals used are immunocompetent but still naïve in their immunological memory (Fig. S5e). Blood samples of the corresponding animals showed no differences in the systemic levels of monocytes or neutrophils between controls and Osr1cKO at 3 dpi (Fig. S5f) excluding systemic effects. Significantly reduced levels of natural killer (NK) cells, but not NKT cells, were found in the injured Osr1cKO muscles (Fig. 4f). Separating macrophages into pro-inflammatory (CD80+) and anti-inflammatory (CD206+) macrophages revealed significantly increased relative numbers of CD80+ macrophages in Osr1cKO muscles, which was accompanied by lower levels of CD206+ macrophages (Fig. 4g, h). While control muscles had a ratio of approximately 1:1 between pro- and anti-inflammatory macrophages, this ratio was shifted to 3:1 in Osr1cKO muscle (Fig. 4h). This finding was confirmed by qPCR examination of whole muscle lysate, which revealed significantly lower expression of *Cd163*, *Cd206* and *Arg1* (anti-inflammatory macrophage markers) in Osr1cKO muscles (Fig. 4i), while expression of *Cd68* and *Cd80* (pro-inflammatory macrophage markers) only tended to be increased (Fig. 4j). Pro-inflammatory macrophages limit the expansion of FAPs through TNFα-induced apoptosis. However, a targeted analysis of transcriptome data showed that genes related to the TNFα signalling pathway are largely downregulated in Osr1cKO FAPs (Fig. S6), suggesting that the cells may be less responsive. In summary, Osr1 depletion does not impair timed leukocyte infiltration into the injured muscle, but affects macrophage polarisation.

### Loss of Osr1 affects MuSCs during regeneration

We next investigated whether the observed delay of muscle regeneration was preceded by altered MuSC function. Immunolabeling for Pax7 showed an approx. 30% decrease in MuSC numbers in the injury area of Osr1cKO muscle at 3 dpi (Fig. 5a). Despite the small lesion size in our model, flow cytometry analysis still detected an approx. 5% reduction of MuSCs levels in Osr1cKO TA muscles at 3 dpi compared to controls (Fig. 5b). Ki67 immunostaining of freshly FACS-isolated MuSCs at 3 and 7 dpi showed an approx. 60% reduction of Ki67+ MuSCs in Osr1cKO animals (Fig. 5c, d). TUNEL staining indicated increased apoptosis of MuSCs in Osr1cKO mice (Fig. 5e).

**Figure 5.**
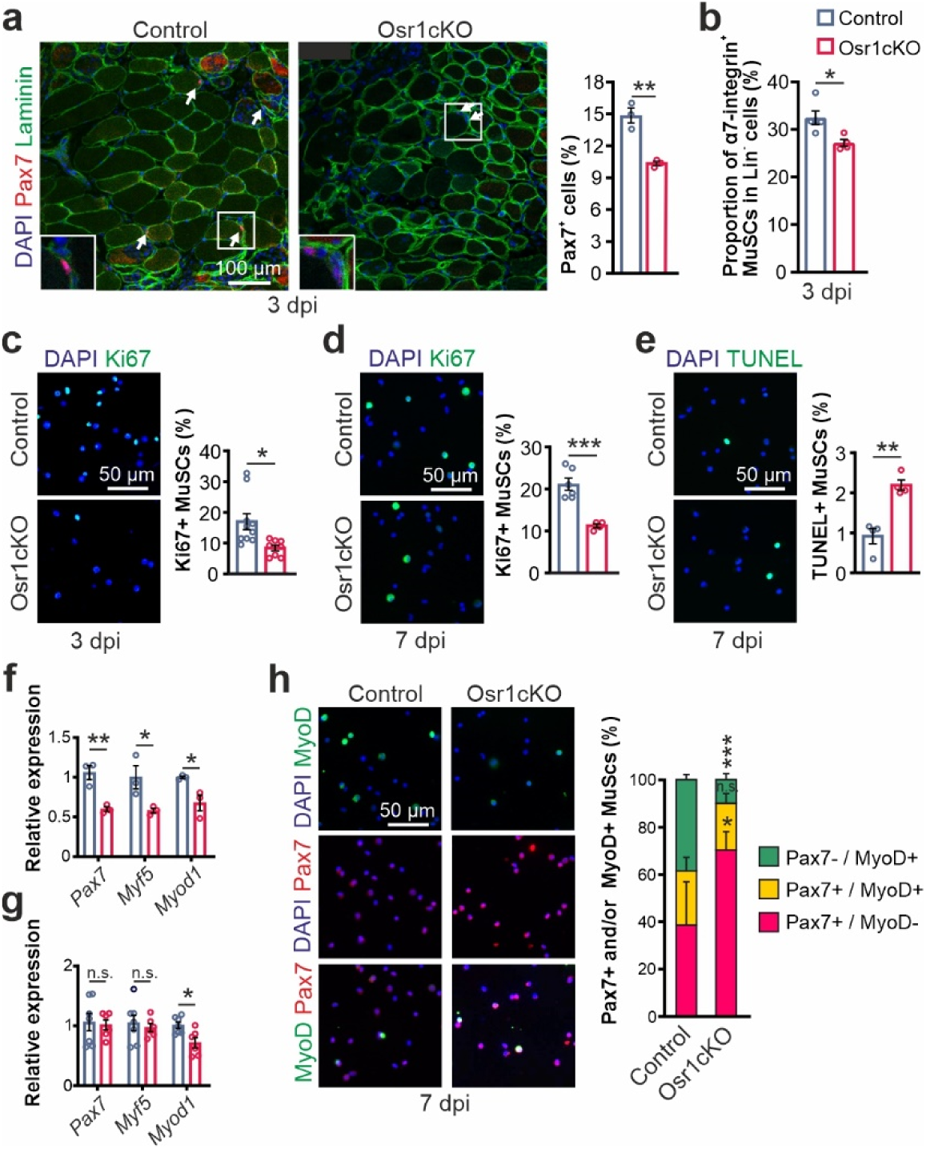
Non-cell autonomous defect of MuSCs in Osr1cKO muscle. **a** Pax7 staining on 3 dpi tissue sections of control and Osr1cKO TA muscle; myofibers are outlined by Laminin, nuclei are stained with DAPI. Quantification of Pax7+ cells in the regenerative area is shown right (n=3 animals per genotype). **b** Flow cytometric analysis of MuSC numbers in Lin-cells at 3 dpi (n=5 for control and n=4 for Osr1cKO). **c, d** Ki67 labeling of MuSCs isolated by FACS at 3 and 7 dpi; representative images left, quantification right (at 3 dpi n=10 for control and n=8 for Osr1cKO, at 7 dpi n=6 for control and n=4 for OsrcKO). **e** TUNEL labeling of MuSCs isolated by FACs at 7 dpi; representative images left, quantification right (n=4). **f, g** RT-qPCR for *Pax7*, *Myf5* and *Myod1* performed on MuSCs isolated by FACs at 3 dpi and 7 dpi (at 3 dpi n=3, at 7 dpi n=7 for control and n=6 for Osr1cKO; each dot represents the mean of three technical replicates from one biological replicate). **h** Immunolabeling for Pax7 and MyoD on MuSCs isolated by FACs at 7 dpi; quantification shown right (n=4). Data are mean ± SEM; *P*-value calculated by two-sided unpaired *t*-test; * p < 0.05, **p < 0.01, ***p < 0.001 N-numbers indicate biological replicates (mice per genotype).

To gain insight into the activation state of MuSCs at 3 and 7 dpi, mRNA expression of *Pax7*, *Myf5*, and *Myod1* was quantified in FACS-sorted MuSCs (Fig. 5f, g). The expression of all three genes was significantly reduced in Osr1cKO MuSCs at 3 dpi (Fig. 5f), indicating a universal defect in the resting and activated states of MuSCs. While *Myf5* and *Pax7* levels were comparable at 7dpi, expression of *Myod1* remained significantly lower at 7 dpi (Fig. 5g). Immunostaining for Pax7 and MyoD on freshly FACS-isolated MuSCs demonstrated a 30% relative increase in Pax7+/MyoD-self renewing cells in the Osr1cKO animals, a similar number of Pax7+/MyoD+ transit amplifying cells, and a 40% reduction in Pax7-/MyoD+ committed cells (Fig. 5h). These results indicate that loss of Osr1 expression in FAPs affects the function of MuSCs in a non-cell autonomous manner and compromises their activation, proliferation and survival, contributing to the delayed muscle regeneration.

### Osr1cKO FAPs affect myogenesis via TGFβ signaling

We next aimed to unravel how Osr1-deficient FAPs may affect MuSC function. FAPs were proposed to promote myogenesis via secreted factors ^6,32^. To test for paracrine effects of Osr1cKO FAPs on myogenesis, we performed indirect transwell co-culture experiments. We used 7 dpi Osr1cKO or control FAPs that were isolated from contralateral muscles of injured animals to achieve an activated “alert” state ^33^, and C2C12 cells (Fig. 6a). Four days after induction of differentiation, fusion of C2C12 cells into myofibers was almost 33% lower in co-cultures with Osr1cKO FAPs compared control FAPs (Fig. 6b). To confirm this result, we used conditioned medium (CM) generated from Osr1cKO or control FAPs to treat C2C12 cells (Fig. 6c). While conditioned medium of control FAPs had no effect on myogenic differentiation of C2C12 cells (Fig. 6d), myogenesis was reduced by approx. 80% in cultures with CM from Osr1cKO FAPs compared to control media (Fig. 6d). These results suggest that Osr1cKO FAPs actively suppress myogenesis via secreted factors. Subsequent search of our transcriptome data for signaling molecules known to negatively affect myogenesis revealed TGFβ pathway-associated GO terms enriched in Osr1cKO FAPs (Fig. S6a, b), and *Tgfb* genes were upregulated (Fig. S7a) in Osr1cKO FAPs. Upregulation of *Tgfb1* was also confirmed in FAPs after *in vitro* recombination (Fig. S7b), suggesting increased *Tgfb1* expression is independent from the injury microenvironment.

**Figure 6.**
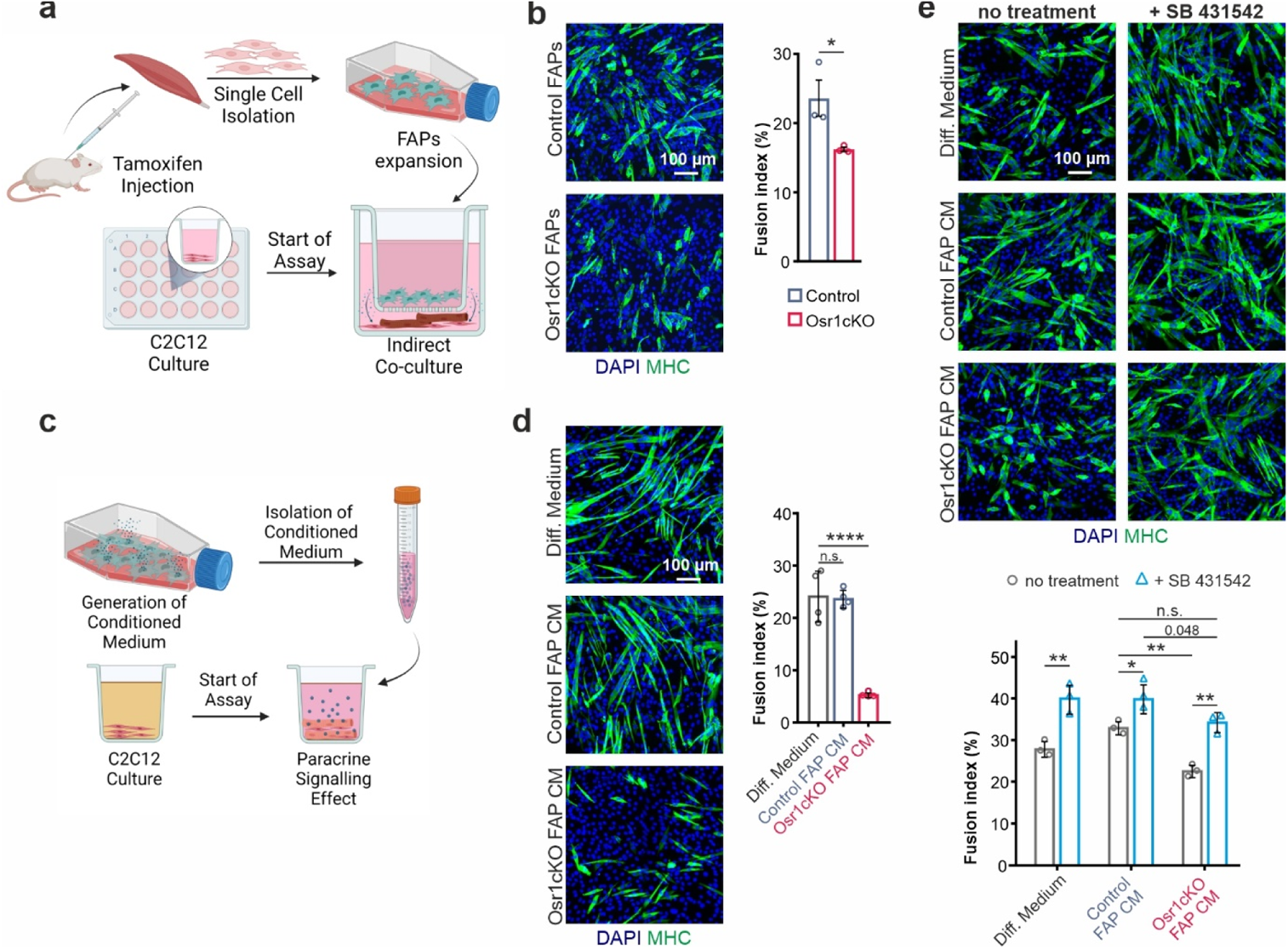
Osr1cKO FAPs inhibit myogenesis via TGFβ signaling. **a** Schematic representation of the transwell assay. **b** Immunolabeling for MHC to detect myotube formation from C2C12 cells co-cultured with control or Osr1cKO FAPs; quantification of fusion index is shown right (n=3). **c** Schematic representation of conditioned medium (CM) assay. **d** Immunolabeling for MHC to detect myotube formation from C2C12 cells in differentiation medium or differentiation medium supplemented with control or Osr1cKO CM; quantification of fusion index is shown right (n=4). **e** Immunolabeling for MHC to detect myotube formation from C2C12 cells in differentiation medium or differentiation medium supplemented with control or Osr1cKO CM, with or without TGFβ pathway inhibitor SB431542; quantification of fusion index is shown below (n=3). Data are mean ± SEM; P-value calculated by two-sided unpaired t-test; * p < 0.05, **p < 0.01, ****p < 0.0001 N-numbers indicate biological replicates (mice per genotype).

TGFβ signaling inhibits myogenic differentiation and myoblast fusion by suppressing *Myod1* and *Myog* expression ^34,35^ and the control of actin cytoskeleton-related genes ^36^. To assess the relevance of TGFβ signaling in CM of Osr1cKO FAPs, TGFβ Type1 receptor kinase inhibitor SB431542 was used. Treatment of C2C12 cells with SB431542 alone reduced phosphorylated SMAD 2/3 levels (Fig. S8a), resulted in elevated myogenin and MHC levels (Fig. S8b), and promoted a nearly 50% higher C2C12 fusion (Fig. 6e), in line with previous observations ^34–36^. Similarly, SB431542 also significantly promoted myogenesis in cultures with CM of control FAPs (Fig. 6e). Importantly, the negative effects of CM from Osr1cKO FAPs on C2C12 differentiation was completely abolished by the addition of SB431542 and fusion rate increased by approx. 40% to the levels of control FAPs CM (Fig. 6e). These results show that Osr1 regulates the paracrine signaling activity of FAPs both *in vivo* and *in vitro*, and loss of Osr1 leads to inhibition of myogenic differentiation via the TGFβ signaling pathway.

### Osr1-deficient FAPs show a pro-fibrogenic shift with altered ECM expression that affects myogenesis

Tissue fibrosis, observed in Osr1cKO muscle, is intimately linked to increased TGFβ signaling ^37^. Upregulation of TGFβ target genes *Tgfbi, Scx*, *Col7a1*, *Loxl2* and *Timp1* ^38–41^ in 7 dpi Osr1cKO FAPs (Fig. S9a) suggested activation of TGFβ signaling. In line, the NIH pathway datasource Bioplanet 2019 identified the pathway “TGFβ regulation of extracellular matrix” enriched in genes upregulated in 3 dpi Osr1cKO FAPs (Fig. S9b). Consistent with the fibrotic phenotype of Osr1cKO FAPs, ECM-coding genes were upregulated at 3 dpi and 7 dpi (Fig. 7a, b). However, ECM-associated GO terms were also found enriched in downregulated genes at 7 dpi (Fig. 3k). Downregulated ECM genes at 7 dpi encoded proteins of the myofiber basal lamina (Fig. 7b), which is in line with the observed insufficient myofiber formation. Conversely, fibrosis-associated structural ECM-components and modifying enzymes were upregulated Osr1cKO FAPs at 7 dpi. These included known fibrotic markers as *Loxl2* or *Timp1* (Fig. 7b).

**Figure 7.**
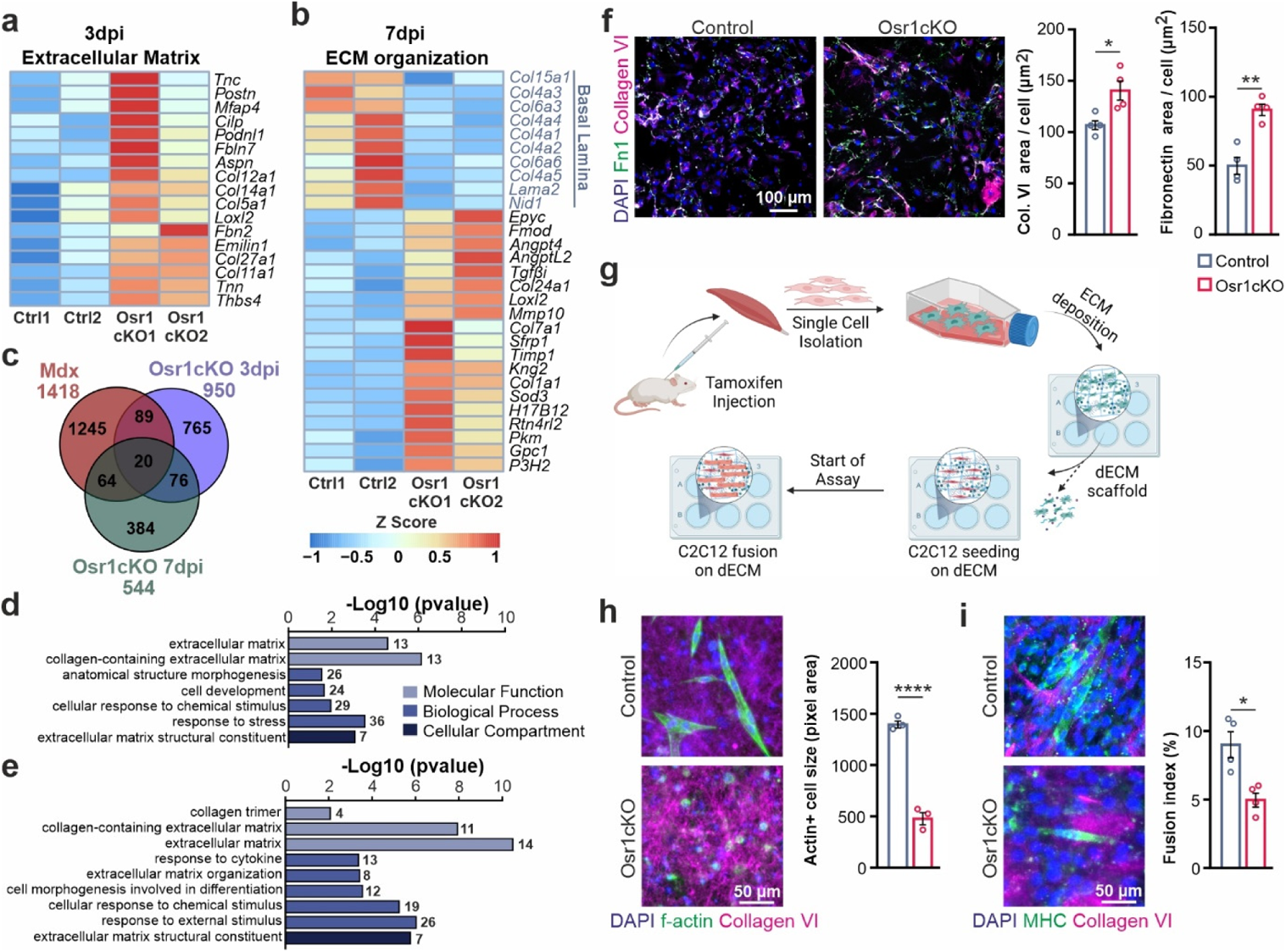
Osr1 deficient FAPs acquire a pro-fibroblastic identity and regulate myogenesis via the ECM. **a, b** Heat maps depicting extracellular matrix (ECM) related genes at 3 dpi and 7 dpi in control or Osr1cKO FAPs. **c** Venn diagram showing overlapping gene deregulation between mdx FAPs and Osr1cKO FAPs at 3 dpi or 7 dpi. **d, e** GO terms analysis of the common deregulated genes between mdx FAPs and the 3 dpi (d) or 7 dpi (e) Osr1cKO FAPs. **f** Immunolabeling for collagen VI and fibronectin (Fn1) on *in vitro* recombined FAPs after 6 days of culture. Quantification of the stained areas shown right (n=4). **g** Schematic depiction of the *in vitro* matrix deposition assay. **h** Immunolabeling for Collagen VI and staining for f-actin using phalloidin on C2C12 cells cultured for two days in differentiation medium on dECM from control or *Osr1 in vitro* recombined cells. Actin staining and its quantification revealed a defect in the morphology of the attached cells on the Osr1cKO dECM. Quantification of f-actin+ cell size shown right (n=3). **i** Immunolabeling for Collagen VI and MHC on C2C12 cells cultured for five days in differentiation medium on dECM from control or *Osr1 in vitro* recombined cells. Quantification of fusion index shown right (n=4). Data are mean ± SEM; P-value calculated by two-sided unpaired t-test; * p < 0.05, **p < 0.01, ****p < 0.0001 N-numbers indicate biological replicates (mice per genotype).

Persistent inflammation and fibrosis are hallmarks of muscular dystrophies including Duchenne muscular dystrophy. In line, comparing our Osr1cKO FAP to mdx FAP transcriptome data ^42^ revealed 173 common DE genes shared between mdx and Osr1cKO FAPs (Fig. 7c). GO analysis of shared DE genes highlighted ECM-related terms as “collagen containing extracellular matrix” between mdx FAPs and 3 dpi (Fig. 7d) as well as 7 dpi (Fig. 7e) Osr1cKO FAPs. Heatmap depiction showed an at least in part common direction of ECM gene deregulation between mdx FAPs and Osr1cKO FAPs (Fig. S9c, d).

To estimate the relative composition of our bulk RNA-seq, we used the deconvolution algorithm MuSiC ^43^ on the annotated single cell dataset of regenerating muscle ^26^. Deconvolution indicated a phenotypic transcriptional shift of Osr1cKO FAPs towards the tenocyte cluster (Fig. S9e) in line with upregulation of tendon- and osteo-chondrogenic transcripts at 3 and 7 dpi (Fig. S9f, g). Remarkably, the “*Osr1* cluster” defined by Oprescu et al. to mainly express genes related with myofiber basal lamina and secreted signaling molecules ^26^ almost completely disappeared in the Osr1cKO FAP population (Fig. S9e). In line, half of the top 51 transcripts from this cluster were downregulated in Osr1cKO FAPs (Fig. S9h). In agreement with increased collagen VI deposition *in vivo* (Fig. S3a), *in vitro* recombined Osr1cKO FAPs showed higher collagen type VI and fibronectin secretion (Fig. 7f) confirming increased fibrogenic differentiation. To investigate the direct effects of altered ECM deposition on myogenesis, FAPs were freshly isolated from contralateral muscles of injured control and Osr1cKO animals. Cells were cultured for 21 days to enable the deposition of a coherent ECM (Fig. 7g, Fig. S10). This *in vitro* formed ECM was subsequently decellularized (dECM) and repopulated with C2C12 myoblast. C2C12 cells cultivated on control dECM for 2 days under differentiation conditions showed a spindle-shaped morphology, while cells on the Osr1cKO dECM remained circular and failed to align and spread (Fig. 7h). Assessing the fusion rate of C2C12 cells on control dECM revealed large multinucleated myotubes after 5 days of culture (Fig. 7i). In contrast, myogenic differentiation of C2C12 cells cultured on Osr1cKO dECM was strongly impaired (Fig. 7i). This indicates that loss of Osr1 in FAPs induces a fibrogenic transcriptional shift in part resembling dystrophic FAPs, resulting aberrant ECM deposition compromising myogenic differentiation.

## Discussion

Our results demonstrate an essential role of the transcription factor Osr1 in the regenerative function of FAPs during muscle healing. Depletion of Osr1 resulted in a skewed inflammatory response, persistent fibrosis, altered mechanical properties of the transient ECM, impaired timely resolution of tissue necrosis, impaired MuSC activation, and ultimately delayed formation and maturation of new muscle fibers.

Regulation of FAP proliferation and elimination is essential for muscle regeneration and prevents fibrosis ^14,21,42,44–48^. Loss of Osr1 reduced FAP pool expansion in a cell-autonomous manner. Pharmacological inhibition of FAP expansion or genetically induced depletion of FAPs resulted in impaired muscle regeneration and fibrosis ^15,16^, in part overlapping our model. However, the amount of FAP cells was only slightly reduced at 3 dpi in the Osr1cKO mice and their levels were comparable to controls at 7 dpi. This suggests that reduction of FAP numbers may have a neglectable contribution to delayed healing in our model, but that loss of Osr1 compromises FAP function comparable to a loss of FAP cells.

Our *in vitro* data show that Osr1-deficient FAPs switch from a beneficial pro-myogenic phenotype to a detrimental state actively suppressing myogenesis. Recent single cell studies have identified distinct FAP subpopulations that either coexist or represent temporally controlled cellular phenotypes during regeneration ^26,49–52^. In addition, tendon progenitors have been defined in muscle as a cell type closely related to FAPs ^49,50^. Loss of Osr1 shifts the phenotype of the FAP population toward a more fibrotic, tendon- or cartilage-like profile, overlapping our previous observations in developmental FAP-like cells ^24^. Moreover, deconvolution analysis indicated a reduction in the representation of a single cell cluster defined by the expression of *Osr1* itself as a hallmark gene ^26^. This suggests that Osr1 is a key regulator of a group of genes defining a specific FAP subtype or state, and prevents FAPs from acquiring alternative fibroblastic states such as tenocyte. Moreover, loss of Osr1 promotes osteo-chondrogenic gene expression in FAPs, reminiscent of osteo-chondrogenic FAP fate in heterotopic ossification ^53,54^.

The *Osr1*+ cluster contains genes encoding proteins of the myofiber basal lamina, but also secreted signaling molecules ^26^, many of which are involved in regulation of immune responses. This suggests a role for the *Osr1*+ FAP population in myofiber maturation and immunomodulation. Skeletal muscle regeneration starts with an inflammatory phase, characterized by immune cell infiltration, locally elevated levels of pro-inflammatory cytokines, and active degeneration of existing muscle fibers ^1,55^. Neutrophils are the first immune cells to arrive in the regenerative region ^56^. Following this, the predominant immune cells are macrophages, which are first activated towards a pro-inflammatory phenotype. Pro-inflammatory macrophages continuously shift their profile to an anti-inflammatory phenotype, resolving inflammation, promoting MuSC differentiation, FAP survival and matrix remodeling ^22,48^. The timely resolution of the acute inflammation is essential for subsequent healing stages, which restores tissue structure and function ^57^. Our data provide first evidence that FAPs directly influence macrophage polarization *in vivo*. While Osr1 deletion did not affect the infiltration of neutrophils and macrophages in the injured region, the transition of the pro- to anti-inflammatory phenotype appeared to be impaired, leading to increased levels of pro-inflammatory relative to anti-inflammatory macrophages. Several of the cytokines downregulated in Osr1cKO FAPs have been involved in macrophage polarization, suggesting direct crosstalk. CCL2 via its receptor CCR2 induces anti-inflammatory polarization ^58^ and inhibits pro-inflammatory cytokine production ^59^, and both CCL8 ^60^ and CCL11 (eotaxin) ^61^ act as anti-inflammatory macrophage recruiters in cancer metastasis. Collectively, these data suggest that Osr1+ FAPs play an important immunomodulatory role in the early phase of healing and that loss of Osr1 expression may lead to a pro-inflammatory environment.

Exacerbated inflammation, in turn, has undesirable consequences for the function of FAPs and MuSCs. For example, increased TNF-α and NF-κB signaling inhibits MuSC differentiation by suppressing *Myod1* expression ^62^. Furthermore, pro-inflammatory macrophages limit FAP expansion via TNFα-mediated apoptosis ^22^. However, increased apoptosis of FAPs was also evident *in vitro*, independent of macrophages. Thus, it is unlikely that skewed macrophage polarization drives increased FAP apoptosis we observed in Osr1cKO muscle. Genes involved in the TNF-α signaling cascade were in fact largely downregulated in Osr1cKO FAPs, suggesting that Osr1-deficient FAPs may be less responsive to TNF-induced apoptosis. This might in turn explain their persistence during later phases of regeneration, causing increased ECM deposition and fibrosis.

FAPs are the major producers of a pro-regenerative transient ECM during regeneration ^50,63^, while an Osr1cKO FAP-derived ECM inhibited myogenesis *in vitro*. Osr1cKO FAPs showed a pronounced shift in ECM gene expression characterized by down-regulation of genes associated with myofiber basal lamina and the upregulation of several ECM molecules that are usually not or only to a minor extent expressed in skeletal muscles. The latter includes the expression of *Col5a1*, *Col11a1*, *Col12a1*, *Tnc* and *Postn*, but also ECM- or signaling-related genes associated with a pro-inflammatory environment, such as Fibromodulin (Fmod) ^64,65^, Angiopoietin-like 4 ^66^, angiopoietin-like 2 ^67,68^ or Secreted Frizzled-related protein 1 (Sfrp1)^69,70^. Our *in vivo* data showed that this altered expression of ECM components coincides with altered mechanical properties of regenerating Osr1cKO muscles. While wild type muscles show a transient increase in tissue stiffness during healing, Osr1cKO muscles showed significant tissue softening. Aberrant ECM deposition and tissue stiffness could be a direct cause for the observed delay in myofiber formation and maturation, as cells sense ECM composition but also rigidity ^71^. Increased matrix stiffness is beneficial for the proliferation and differentiation of MuSCs ^63,72–74^. Conversely, decreased stiffness is associated with the formation of fibrotic scar tissue during impaired muscle regeneration ^63^, various pathologies^28,75–77^, or aging^78,79^. This altogether suggests that Osr1cKO FAPs produce an inadequate fibrotic replacement matrix during the early phase of regeneration directly interfering with myogenesis.

Impaired muscle regeneration and fibrosis in disease or aging are closely linked do increased TGFβ signaling ^30,80–82^. Moreover, in muscle dystrophy, asynchronous waves of inflammation lead to a chronic TGFβ rich environment ^48,83,84^. Fibrosis in mdx mice can be exacerbated by injection of TGFβ, and application of TGFβ concomitant to injury in wild type mice resulted in accumulation of fibrotic ECM ^85^ comparable to our model. Osr1-deficient FAPs showed increased expression of *Tgfb1*, and we show that the myogenesis-inhibiting effect of Osr1cKO FAPs can be counteracted by blocking TGFβ signaling with a receptor kinase inhibitor. This is in line with previous reports showing a similar anti-myogenic effect of enhanced TGFβ signaling in muscular dystrophies or during regeneration ^36,86,87^. Of note, increased TGFβ signaling in the Osr1cKO muscle microenvironment may at the same time impair myogenesis, and also in an autocrine fashion promote the pro-fibrogenic phenotype of FAPs contributing to persistent fibrosis. This is supported by the enrichment of and of TGFβ pathway-related genes and upregulation of TGFβ pathway downstream targets in the transcriptome of Osr1cKO FAPs.

In conclusion, our studies show that FAPs are key mediators of an intricate balance coordinating inflammation and regenerative myogenesis, and that Osr1 is an essential transcriptional regulator of the pro-regenerative FAP phenotype.

## Methods

### Mice

Mouse lines were maintained in an enclosed, pathogen-free facility. All experiments were performed in accordance with the European Union legislation for the protection of animals used for scientific purposes, and approved by the Landesamt für Gesundheit und Soziales Berlin under license numbers ZH120, G0114/14 and G0198/19. CAGG-CreER mice were described before^88^. Osr1^flox^ mice were derived from an *Osr1* multifunctional allele (Osr1^MFA^; ^25^) by crossing with ubiquitous flippase mice ^89^.

### Muscle Injury and tamoxifen administration

The tibialis anterior muscle of 4-6 months old mice was injured using the freeze-pierce method. Mice were anaesthetized using isofluoran (Univentor 410 anesthesia unit), or by intraperitoneal injection of 10% (v/v) ketamine / 2% (v/v) xylazine (Rompun® 2%) in sterile PBS (5 μl / g body weight), while animals were on a 37°C heating plate. After that, the skin above the tibialis anterior muscle was opened and the muscle was pierced five times using a syringe needle pre-cooled in liquid nitrogen. On the day before injury until 2 days post injury, analgesia was performed using Carprofen. Tamoxifen (Sigma-Aldrich) was dissolved in a mixture of 90% sunflower oil and 10% ethanol. Animals were injected i. p. with 150µl of a 20 mg/ml Tamoxifen stock on the day of injury (prior to injury) and on the two following days. As control CAGG-CreER^neg^;Osr1^flox/flox^ or CAGG-CreER^neg^;Osr1^flox/+^ animals were used for histological analysis and stiffness measurements, while CAGG-CreER^+^;Osr1^flox/+^ animals were used as control for experiments involving FACS of Osr1-mGFP+ FAPs (transcriptome analysis and *in vitro* assays).

### RNA extraction and RT-qPCR

Total RNA isolation was performed using the Direct-zol RNA Microprep kit (for in-vitro experiments and FACS sorted primary cells) and the Direct-zol RNA Miniprep kit (for whole muscle lysate). Reverse transcription for cDNA synthesis was performed using M-MuLV Reverse Transcriptase (Enzymatics, Qiagen) and RNAse Inhibitor (Biotechrabbit). RT-qPCR was performed in a 384-well plate, in a 12 µl mixture consisting of 6 µl Blue SYBR Green mix (Biozym) or GOTaq qPCR Master Mix (Promega), 2 µl primer pairs (2.5 µM each) and 5 µl template of cDNA. RT qPCR was performed in three technical replicates from each of at least three biological replicates (representing individual animals or cells derived from one animal). Analysis was performed using the ABI Prism HT 7900 real time PCR detection system (Applied Biosystems) equipped with SDS software version 2.4 (ThermoFisher Scientific) or the QuantStudio 7 (Applied Biosystems) with the software version 1.3 (ThermoFischer Scientific). The mean relative values of the three technical replicates were normalized to the values of GAPDH, which was used as a housekeeping gene. To calculate the relative expression level of the respective gene, the double delta Ct (ΔΔCt) method was used. All primers used were purchase from Eurofins Scientific and are listed in Table S1.

### Tissue preparation and histology

Directly upon dissection, muscle was embedded in 6% (w/v) gum tragacanth (Sigma-Aldrich) dissolved in H_2_O, and snap frozen in ice-cold isopentane (precooled in liquid nitrogen, −160°C). Muscle tissue was sectioned at 10 µm thickness onto Superfrost Plus slides (Thermo Scientific) and stored in −80°C until use. Hematoxylin and Eosin (Thermo Fisher) (H&E) staining and picrosirius red staining were performed according to standard procedures. Stained samples from n=3 mice per genotype per timepoint (3, 5, 10,17 and 28 dpi) were imaged in total using a Leica brightfield microscope equipped with an automated XY scanning stage. The area size of the collagenous matrix deposition was quantified in Image J from the picrosirius red staining and was further normalized to the size of the total area size of the whole regenerative region.

### Immunolabelling

Prior to antibody labelling, sections were allowed to reach room temperature (RT) slowly, slides were immersed in PBS for 5 min. and fixed in 4% PFA for 10 min. at RT. Slides were shortly washed in PBS and then permeabilized with 0,4% (v/v) Triton X-100 (Sigma Aldrich) in phosphate buffer (PBS) for 10 min. Sections were blocked with 5% bovine serum albumin (Sigma Aldrich) in 0,1% Triton X-100 in PBS for 1 hour at RT. Primary antibodies were diluted in 5% BSA in 0,1% Triton X-100 in PBS and incubated overnight at 4°C, the following day slides were washed three times in PBS for 5 min. Secondary antibodies were diluted in 5% BSA in PBS and incubated for 1 hour at room temperature, followed by three short washes in PBS for 5 min. Nuclei were stained with 5 µg / µl 4′,6-diamidino-2-phenylindole (DAPI; Invitrogen) and slides were mounted with FluoromountG (SouthernBiotech).

For MHC3 immunostaining, antigen retrieval was performed upon fixation. Sections were treated in chilled methanol, kept in −20°C for 6 min, and subsequently washed in PBS. Then, slides were immersed in 1 mM Ethylendiaminetetraacetic acid (Roth) at 95°C for 10 min. The slides were left at RT for 30 min and washed in PBS. Blocking and antibody staining was performed as described above.

For Pax7 immunostaining, upon fixation with 4% PFA, slides were permeabilized in chilled methanol for 6 min at RT. Epitope retrieval was performed in boiling 1mM citric acid pH 6 for 2 min and 30 sec in a microwave. Slides cooled to RT, washed in PBS and blocked in 5% BSA IgG-free (Jackson Immuno Research) in PBS. To decrease background, sections were incubated with anti-mouse IgG Fab fragments (Jackson Immuno Research) diluted in PBS 1:100 for 30 min at RT. Primary and secondary antibody were diluted in 5% BSA IgG free and staining was performed as described above. A list of primary and secondary antibodies is provided in Tables S2 and S3.

For cell immunostaining, isolated cells were added on the coated coverslips (1 hour at RT in Poly-L-lysine (Milipore) diluted 1:100 in water) and allowed to adhere for 1 hour at 4°C. Then, cells were fixed in 4% PFA for 15 min at RT and washed shortly in PBS. Samples were permeabilized in 0,4% (v/v) Triton X-100 (Sigma Aldrich) in phosphate buffer (PBS) for 10 min. Primary and secondary antibody labelling followed as described above. Cytospun collected cells were stained following the same procedure. TUNEL staining on cytospun and cultured cells was performed using the DeadEnd^TM^ kit (Promega) according to the manufacturer’s instructions.

### Cell sorting of FAPs and MuSCs by FACS

FAP and MuSC isolation from muscle was performed as described in ^25^. Briefly, TA muscle was isolated and roughly minced with a small scissor in high-glucose DMEM medium (PAN Biotech) containing 10% fetal bovine serum (PAN biotech), 1% Penicillin Steptomycin (P/S) solution (PAN Biotech, 10.000 U/ml) and 2,5 mg/ml Collagenase A (Roche) for 45 min at 37°C with gentle shaking. Muscle lysates were further digested with 2 IU/ml of Dispase II (Sigma Aldrich) diluted in minimum amount of PBS for 30 min. To stop the enzymatic digestion, 10% FCS DMEM was added to the sample and subsequently the sample was passed ten times through a 20G syringe needle. Cells were filtered first through a 70 µm and then through a 40 µm cell strainer (Fischer Scientific) and collected by centrifugation at 400xg for 10 min. Cells were resuspended in filtered “FACS buffer” containing 1% BSA and 2 mM EDTA, labeled with the following antibodies: anti-CD45-APC, anti-Cd31-APC, anti-TER119-APC, anti-a7-integrin-PE, and anti-Ly6A/e-APC-Cy7 (Table S4) for 30 min on ice and washed three times with FACS buffer prior to sorting. Propidium iodide was used as a viability dye. Cell sorting and analysis was performed on a BD FACSAriaII SORP (BD Biosciences). Single-stained and fluorescence minus one controls were used for setting the sorting gates. Data were collected using the BD FACSDiva software version 8.0.1.

### Immune cell flow cytometry analysis

Muscles were cut into small pieces and transferred to a gentleMACS C tube (Miltenyi Biotec, Bergisch Gladbach, Germany) containing TrypLE Express Enzyme (ThermoFisher, Waltham, MA, USA) in a 37°C water bath for 5 min. The sample was run on a gentleMACS Dissociator (Miltenyi Biotec, Bergisch Gladbach, Germany) using the predefined program for murine muscle dissociation. The tube was transferred back to the 37°C water bath for another 5 min and the dissociation program was repeated. The dissociated tissue was then filtered through a 40µm nylon mesh (ThermoFisher, Waltham, MA, USA) and thoroughly washed with PBS. Cells were counted and incubated with live-dead stain (LIVE/DEAD Fixable Blue for UV excitation, ThermoFisher, Waltham, MA, USA) at 4°C. Washing steps were performed using PBS supplemented with 0,5 % w/v bovine serum albumin and 0,1 % sodium acid (both Sigma-Aldrich, St. Louis, MO, USA). Surface marker incubation was performed before intracellular staining at 4°C. Intracellular staining for epitopes was achieved using the fixation buffer and intracellular permeabilization buffer kit (BioLegend, San Diego, CA, USA). Antibodies are listed in Table S4. Flow analysis was run on a CytoFlex LX system (BeckmanCoulter, Brea, CA, USA) and population gating and tSNE analysis were performed with FlowJo (BD Biosciences, Franklin Lakes, NJ, USA). The gating strategy was adapted as previously published ^90^. Outliers were identified with the ROUT method and excluded from the analysis.

### Isolation of adherent fibroblasts for *in vitro* FAP culture

Adherent connective tissue fibroblasts that are phenotypically identical to FAPs were isolated from skeletal muscle in essence as described before ^31^, thus we refer to these cells as FAPs. For generation of conditioned medium, co-culture assays and for the decellularization assays, cells were isolated from contralateral hindlimbs of injured 7 dpi animals. This was done to reduce animal usage and to achieve a pre-activated “alert” state of FAPs ^25,33^. For cell isolation, briefly, the TA, gastrocnemius and the quadriceps muscles were carefully isolated and cut with a scissor. Tissue digestion was performed as for flow cytometry. After centrifugation, cells were resuspended in DMEM with 10% FCS, placed in a 10 cm dish and allowed to attach for 90 min at 37°C. Supernatant was removed, containing non-attaching cells, adherent cells were washed once with PBS and fresh DMEM with 10% FCS was added (Passage 0). After 3-4 days of growth, cells were trypsinized, counted using the automated LUNA^TM^ cell counter (Logos Biosystem), and immediately used for the respective assay (Passage 1). For every experiment, freshly isolated FAPs (Passage 1) were used.

### In-vitro recombination of the *Osr1* locus

0.5 µM of 4-hydroxytamoxifen (4-OHT) was added to the medium of passage 0 control or Osr1cKO FAPs. 4-OHT treatment was repeated on day 1 and day 2 of culture, without discarding the old medium. The efficiency of the recombination was measured via genomic qPCR for the exon 2 of *Osr1* and by RT-qPCR. For differentiation assays, cells were trypsinized upon 50-60% confluency to avoid spontaneous differentiation. Approximately 25.000 FAPs were seeded on coverslips in 24 well plates and cultured in DMEM 10% FCS for 6 days. Fresh medium was added every 2 days. Cells were fixed with 4% PFA and immunolabeled as described above

### Indirect co-culture of FAPs and C2C12

For transwell assays, 10.000 *in vitro* recombined control or Osr1cKO FAPs were seeded in the insert and cultured in transwell plates for 14 days in proliferation medium (high glucose DMEM, 10% FBS, 1% P/S). Upon confluency, 100.000 C2C12 cells per well were seeded to the chambers and allowed to attach for 4 hours. Proliferation medium was then removed and replaced with differentiation medium (high glucose DMEM, 2% horse serum (PAN biotech), 1% P/S). After 4 days of differentiation, cells were fixed for immunolabeling. To analyze C2C12 fusion, four different areas per sample were analyzed and the number of nuclei in MyHC+ fibers was normalized to the total number of nuclei.

### Conditioned medium assays

30.000 *in vitro* recombined control or Osr1cKO FAPs were seeded on 24-well plates and expanded in DMEM 10% FCS and 1% P/S until they reached confluency. Cells were briefly washed with PBS, then 300µl of DMEM 1% P/S without FCS was added. After 24h conditioned medium (CM) was collected, spun for 15min in 4°C at 3.000 rcf and supernatant was isolated. CM was stored in −80°C and thawed on ice before use.

For CM treatment, 50.000 C2C12 cells were seeded on coverslips in 24-well plates and kept in proliferation medium (high glucose DMEM, 10% FCS, 1% P/S) for 3 days. Conditioned medium (500 µl) was supplemented with 2% HS and used as differentiation medium for the C2C12 cells. As positive control, fresh differentiation medium (high glucose DMEM, 2% horse serum, 1% P/S) was used. New conditioned medium was added on cells on day 2, and on day 4 cells were fixed with 4% PFA and immunolabeled as described above. Fusion index was calculated as the ratio of nuclei in MHC positive fibers versus the total number of nuclei. Four different areas per sample were imaged and quantified.

For SB431542 treatment, 20.000 C2C12 cells per well were seeded on an 18-well IBIDI μ slide in proliferation medium. On the following day, cells were starved for 5h in high glucose DMEM with 0% FCS and then cells were pre-treated for 1h with 0.5 μM SB431542 (Selleck chemicals). SB431542 medium was aspirated, cells were briefly washed with PBS. Then, 80µl of conditioned medium supplemented with 2% HS and 0.5 μM SB431542 was added to the cells. On day 3 of differentiation, cells were fixed and immunolabeled as described above. Each chamber of the slide was completely imaged, fusion index was quantified as described above.

### Decellularization and dECM assays

To generate the dECM scaffolds, 30.000 *in vitro* recombined control or Osr1cKO FAPs were seeded on 0.1% gelatin (Roth) coated coverslips in 24-well plates. Cells were cultured in matrix medium consisting of high glucose DMEM, 10% FCS, 1% P/S and 50 µM of ascorbic acid. Matrix medium was changed every 2-3 days. Cells were cultured for approximately two weeks until a visible thin layer of matrix formed, after which cells were removed using the prewarmed 0,5% Triton-X-100 and 20 mM NH_4_OH in PBS. Afterwards the three-dimensional matrices were gently washed with PBS and 100.000 C2C12 were seeded in differentiation medium (high glucose DMEM, 2% horse serum (Pan Biotech), 1% P/S) for 2 or for 5 days. Half of the medium was then aspirated and 4% PFA was added for 15 min. at RT. Attached cells and ECM scaffolds were immunolabeled as described above.

### Nanoindentation

The surface elasticity of 15 µm sections from injured TA muscles was measured using the Piuma nanoindenter (Optics11life). The system was calibrated for Young’s modulus measurement, approaching the slide surface and performing an initial wavelength scan. The cantilevers used had a tip radius in the range of 9.5 µm and stiffness of 0.52 N/m. Tissues were immersed in deionized water for 10 min prior to the measurements. Matrix scans of 10X10 were performed setting the step size of the cantilever to 15µm. The data were acquired using the Piuma software.

### RNA sequencing

GFP+ FAPs were isolated via FACS from injured muscles from 4 Controls and 4 Osr1 cKO animals at 3 dpi and from 8 Controls and 8 Osr1cKO animals at 7 dpi. 2 samples each at 3 dpi, and 4 samples each at 7 dpi were pooled and served as a biological replicate. RNA was isolated using the Micro RNA Kit (Zymo Research), the RNA concentration was measured using a QubitFluorometer (Invitrogen), and the quality of the RNA yield was measured with the Bioanalyzer 2100 (Agilent). After quality control using Agilent’s Bioanalyzer sequencing libraries were prepared from 100ng of total RNA per sample following Roche’s stranded “KAPA RNA HyperPrep” library preparation protocol for dual indexed Illumina libraries: First the polyA-RNA fraction was enriched using oligo-dT-probed paramagnetic beads. Enriched RNA was heat-fragmented and subjected to first strand synthesis using random priming. The second strand was synthesized incorporating dUTP instead of dTTP to preserve strand information. After A-tailing Illumina sequencing compatible unique dual index adapters were ligated. Following bead-based clean-up steps the libraries were amplified using 11-12 cycles of PCR. Library quality and size was checked with qBit, Agilent Bioanalyzer and qPCR. Sequencing was carried out on an Illumina HiSeq 4000 system in PE75bp mode and on NovaSeq4000 in PE100bp mode, respectively. Read mapping to the mouse genome (mm10) was performed using STAR in the Galaxy Europe platform, and differential gene expression analysis was performed using DESeq2. Genes were considered as being differentially expressed if the fold-change of KO vs Control was greater than 1.2, if the p-value was below 0.05 for the 3 dpi samples and if the Benjamini-Hochberg adjusted p-value (padj) was below 0.1 for the 7 dpi samples. Transcripts per million (TPM) abundances were calculated from the mean normalized fragment counts given by DESeq2 for all samples. Gene ontology and pathway analysis was performed using the functional annotation tools Enrichr ^91^ and g:Profiler^92^.

### Immunoblotting

Protein isolation was performed upon cell sample homogenization using RIPA buffer (50 mM Tris-Hcl, pH 8.0; 150 mM Nacl; 1% NP-40; 0.5% Sodium deoxycholate; 0.1% SDS). Protein concentration was determined using the Pierce BCA Protein Assay Kit (Thermo Fischer #23225). Total protein was loaded and separated in SDS-page gels and then transferred to PVDF membrane (GE Healthcare). For blocking, 5% BSA in TBST was applied on the membrane for 1h at RT. Primary antibodies were diluted in the blocking buffer and were incubated overnight at 4°C. Membranes were washed three times in PBST, and then incubated with HRP-conjugated secondary antibodies diluted in PBS for 1h at RT. Antibodies are listed in Table S2. The Fusion X spectra gel documentation system (Vilber) was used for image acquisition (FUSION FX software).

### Data Availability

All sequencing data will be made available via GEO upon publication.

## Acknowledgements

We thank the animal facility of the Max Planck Institute for Molecular Genetics, Berlin for expert support; Carmen Birchmeier for providing antibodies; Heiner Schrewe for providing CAGG-CreERt mice. Schematic images were generated using BioRender®. GK was supported by the Berlin-Brandenburg School for Regenerative Therapies (BSRT). GK and SS were funded by the Einstein Foundation Berlin via the Einstein Center for Regenerative Therapies ECRT, and by the Sonnenfeld Stiftung Berlin. SG was funded by the German Federal Ministry of Education and Research (BMBF, 031L0234B), European Union’s Horizon 2020 research and innovation program (Grant No. 779293), and German Research Foundation (DFG; Research Group 2165, GE2512/2-2). SG and PK were funded by the DFG Collaborative Research Center 1444.

## Author contributions

Conceptualization: SS. Supervision: SS, SG, PK, THQ. GK performed the majority of experiments and data collection. THQ, CHB, SPK, VU, PVG performed additional data collection. CGT supervised FACS experiments. CHB performed immune cell profiling experiments. SB and BT performed mRNA-Sequencing. WJ and FLG performed deconvolution analysis of scRNA-Seq data. ANE designed the Osr1-MFA allele. Formal data analysis and interpretation were performed by GK, CHB, PVG, FLG, PK, SG and SS. Writing – Original Draft: SS, GK. Writing – Review & Editing: SS, SG, PK, FLG, ANE.

## Materials & Correspondence

Correspondence and material requests should be sent to S.S.

## Supplementary Materials

### Supplementary Figures

**Figure S1.**
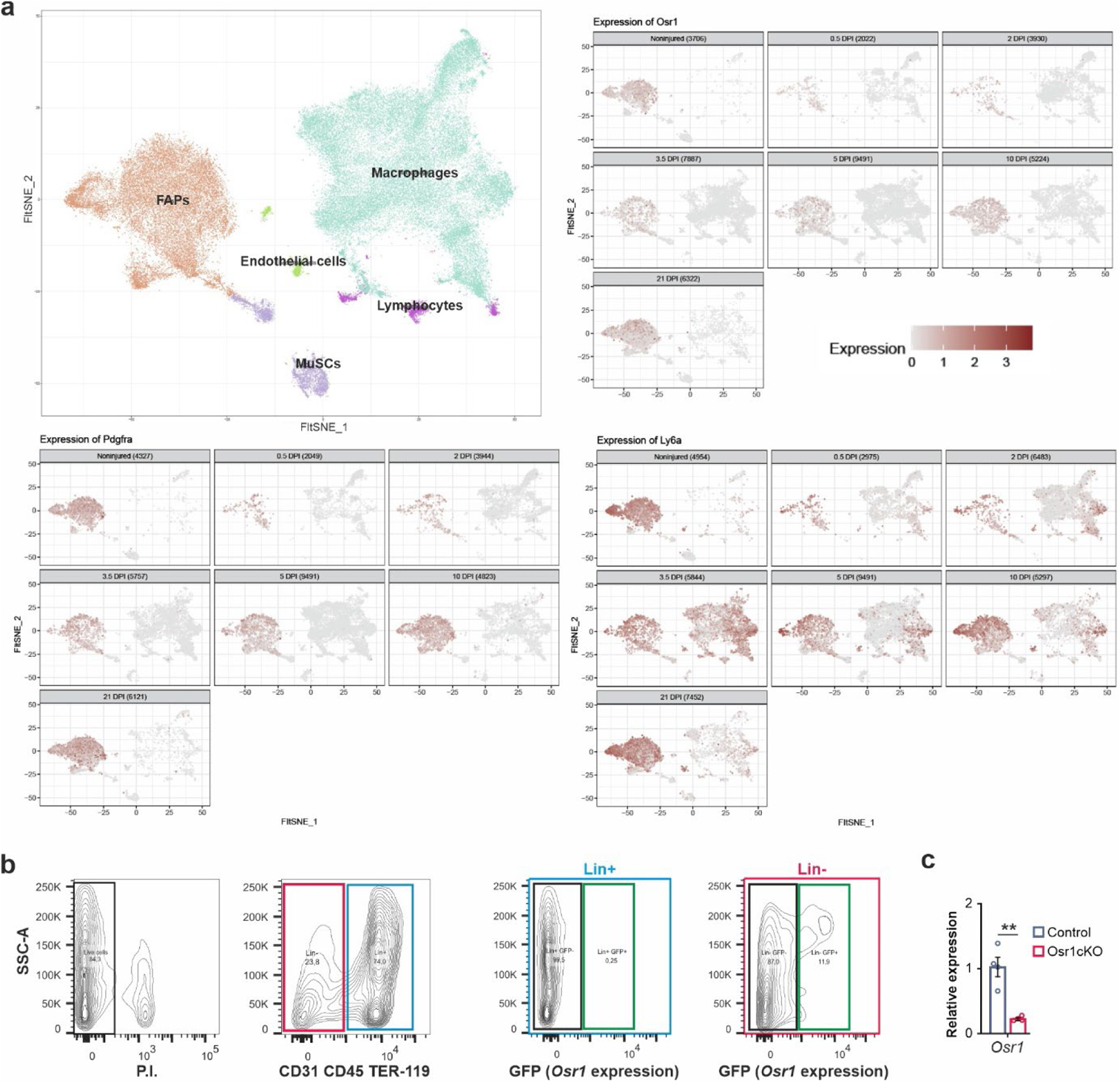
Efficacy of Osr1 conditional inactivation and specificity of Osr1 expression. **a** Analysis of *Osr1* expression in comparison to *Pdgfra* and *Ly6a* (Sca-1) in single cell data from Oprescu et al. of uninjured muscle and of indicated time points post injury. Cluster annotation shown in top left panel. **b** FACS sorting strategy for cells expressing GFP from the recombined Osr1 locus from 3 dpi muscle. Note no GFP+ cells were detected in Lin+ hematopoietic and endothelial cells. **c** RT-qPCR analysis of *Osr1* mRNA expression in whole muscle RNA of control and Osr1cKO mice at 3 dpi *(n=4)*. Data are mean ± SEM; P-value calculated by two-sided unpaired t-test; **p < 0.01. N-numbers indicate biological replicates (mice per genotype).

**Figure S2.**
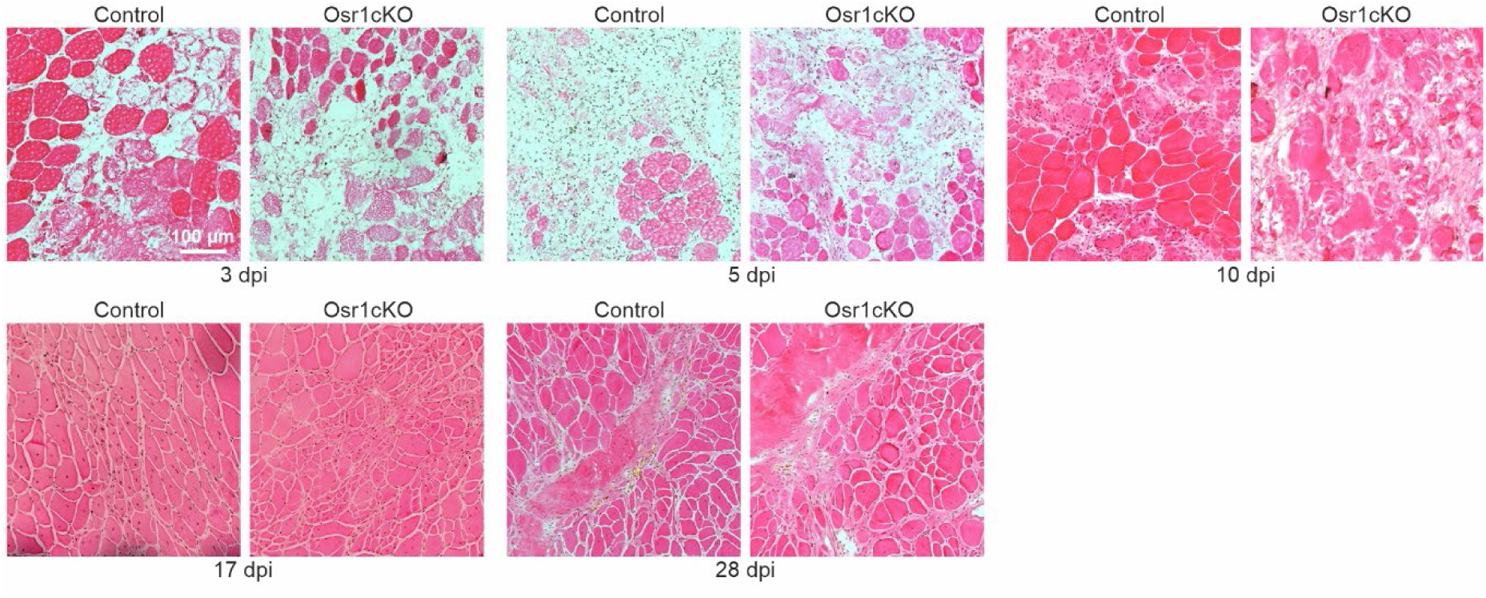
Delayed muscle regeneration and fibrotic appearance of Osr1cKO regenerating muscle. Hematoxylin and eosin staining of control and Osr1cKO muscle sections at indicated days post injury (dpi).

**Figure S3.**
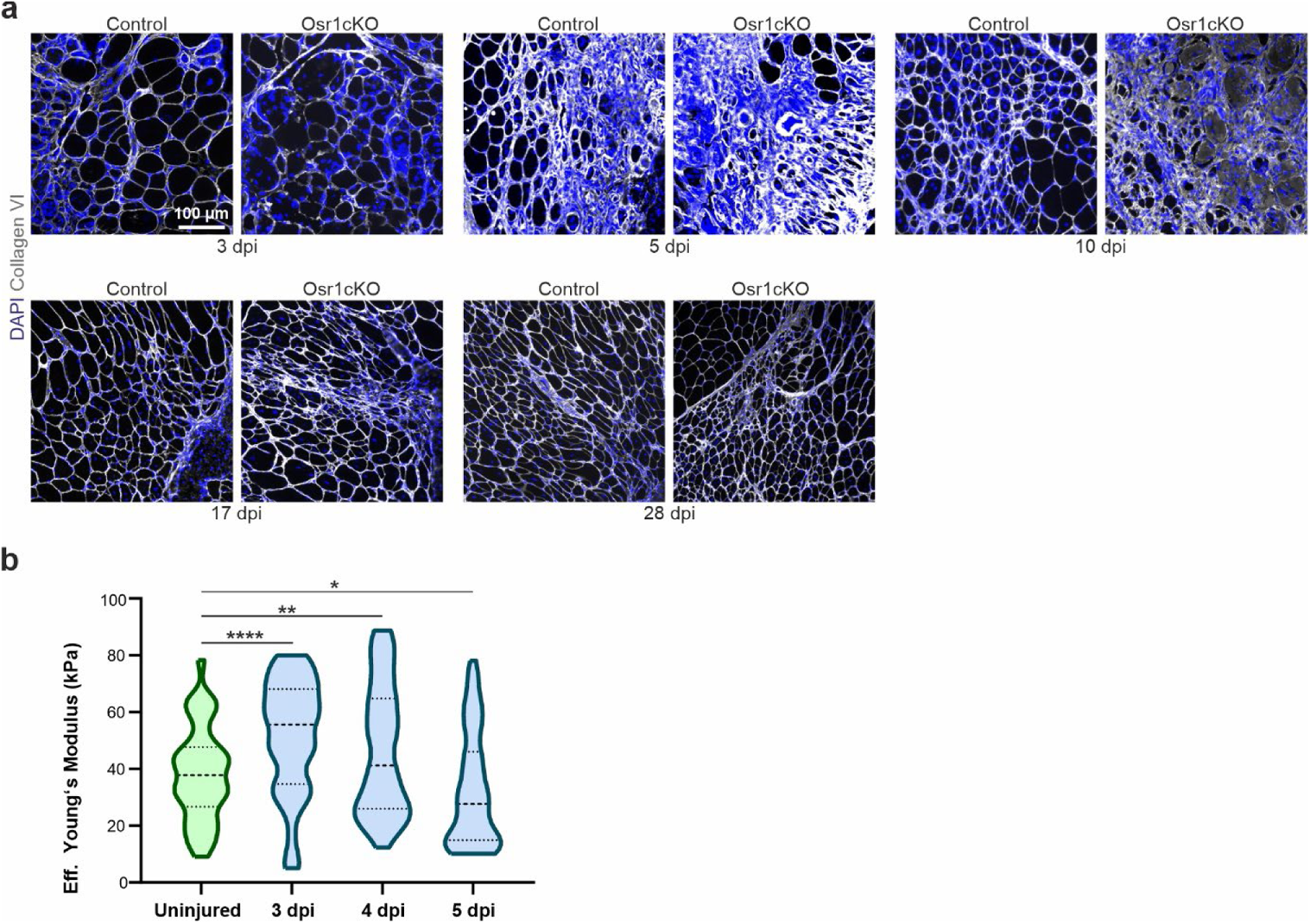
Fibrotic appearance of Osr1cKO regenerating muscle. **a** Immunolabeling for Collagen VI on control and Osr1cKO muscle sections at indicated dpi. **b** Tissue stiffness measurement assessed by nanoindentation of wild type regenerating muscle tissue sections at indicated dpi (n=3). Data are mean ± SEM; P-value calculated by two-sided unpaired t-test; * p < 0.05, **p < 0.01, ****p < 0.0001. N-numbers indicate biological replicates (mice per genotype).

**Figure S4.**
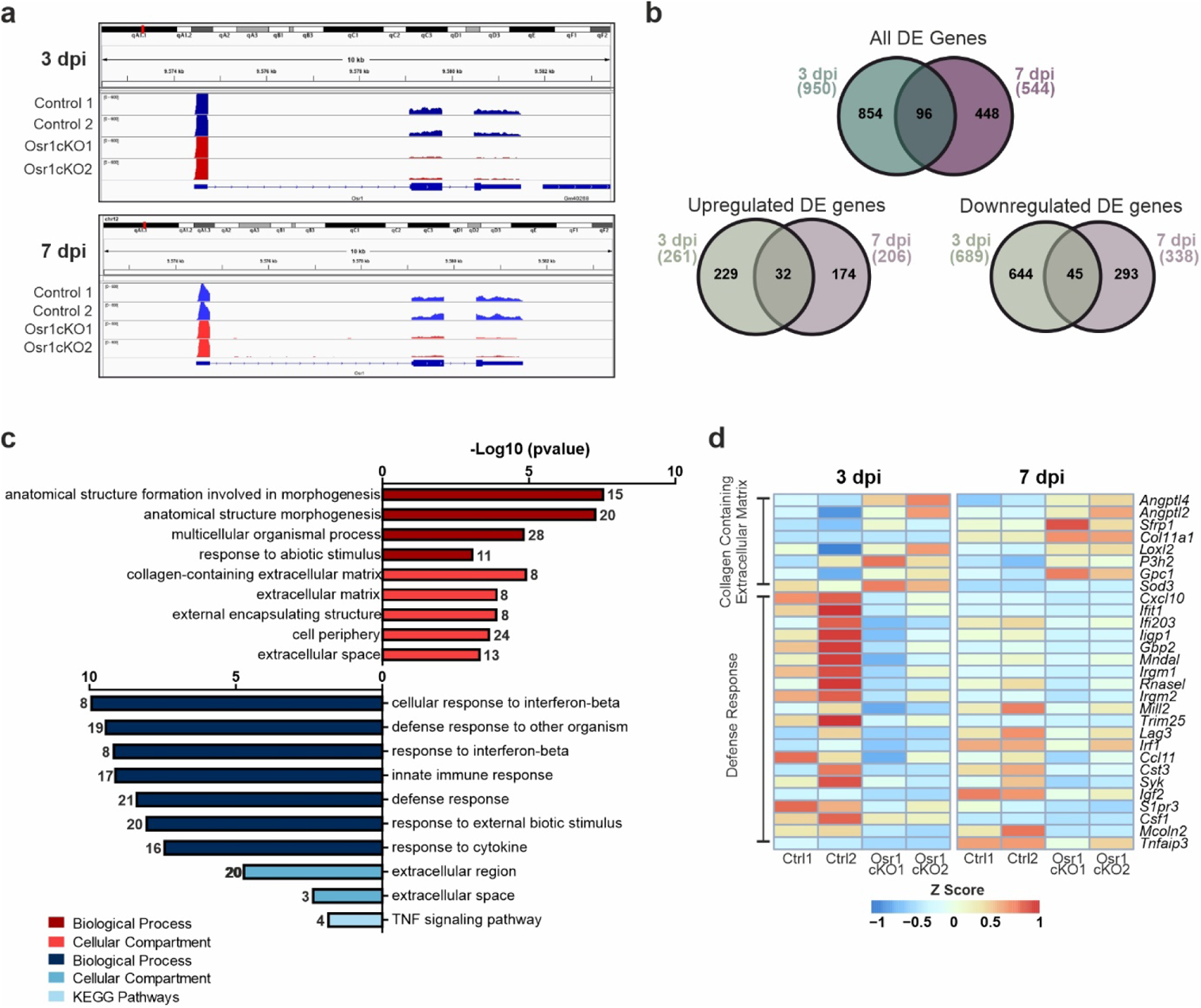
Supplementary transcriptome analysis of Osr1cKO FAPs. **a** Genome browser view of the *Osr1* locus with RNA seq data from 3 and 7 dpi. Note efficient recombination indicated by strongly reduced exon 2 and 3 reads. **b** Venn diagram depicting common regulated genes between the 3 and the 7 dpi Osr1cKO FAPs. **c** GO term analysis of genes commonly upregulated (top) and downregulated (bottom) in Osr1cKO FAPs at 3 and 7 dpi. **d** Heat maps showing genes belonging to the GO terms “Collagen-containing extracellular matrix” and “Defense response” at 3 and 7 dpi showing continuous deregulation.

**Figure S5.**
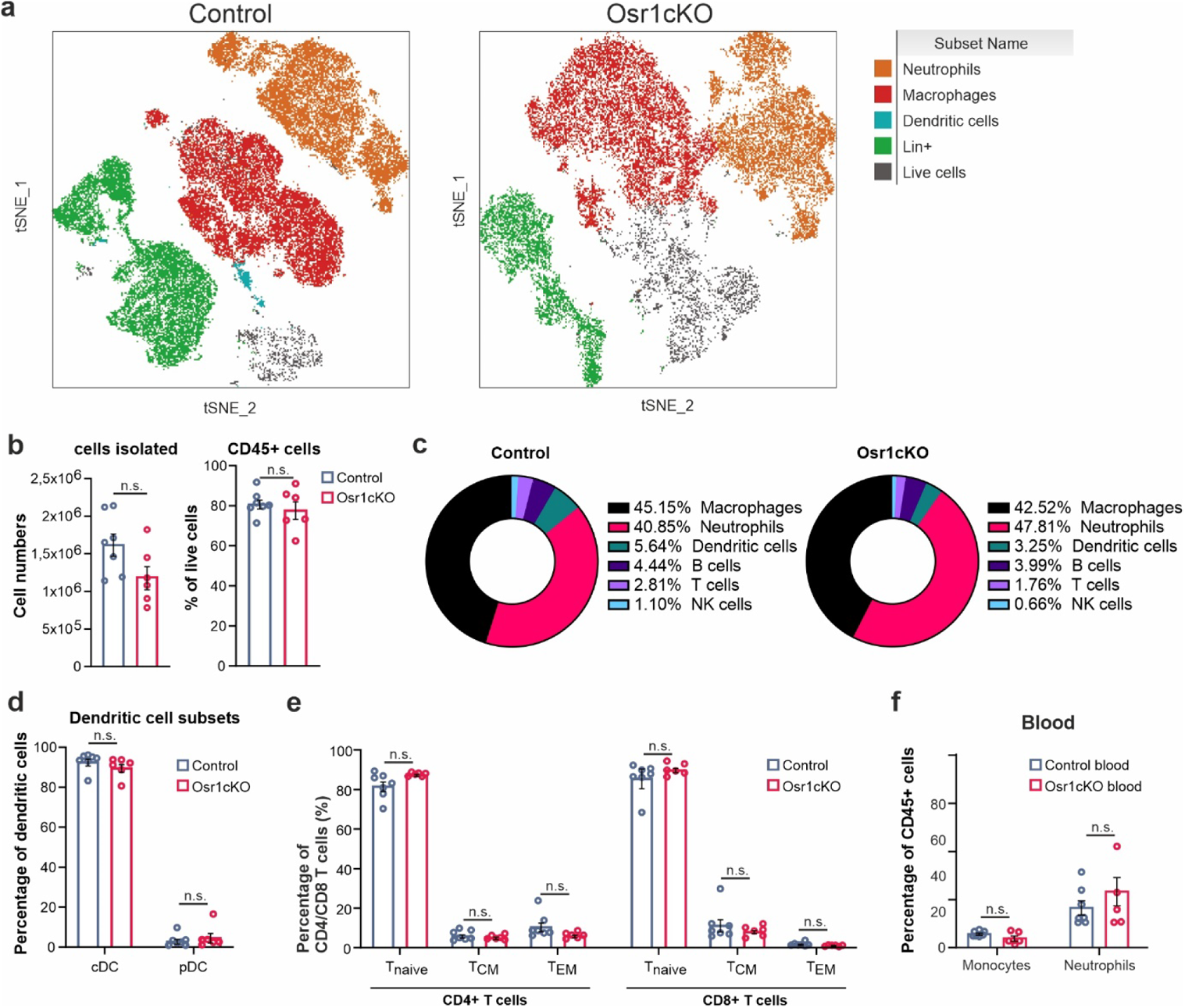
Supplementary data immune cell profiling at 3 dpi. **a** tSNE plot depiction of immune cell populations identified in 3 dpi control or Osr1cKO muscle. Lin+ is defined CD3+, CD19+, CD335+. **b** Flow cytometry quantification of all cells analyzed and percentage of CD45+ cells in 3 dpi control or Osr1cKO muscle. **c** Percentages of immune cell populations identified in control and Osr1cKO muscle. **d** Flow cytometry quantification of Dendritic cell subsets in 3 dpi control or Osr1cKO muscle. **e** Flow cytometry quantification of T-cell cell subsets in 3 dpi control or Osr1cKO muscle. **f** Flow cytometry quantification of macrophages and neutrophils in blood of control and Osr1cKO mice at 3 dpi. In d,c,d and e n=6 for control and n=7 for Osr1 cKO. Data are mean ± SEM; P-value calculated by Mann-Whitney test; N-numbers indicate biological replicates (mice per genotype).

**Figure S6.**
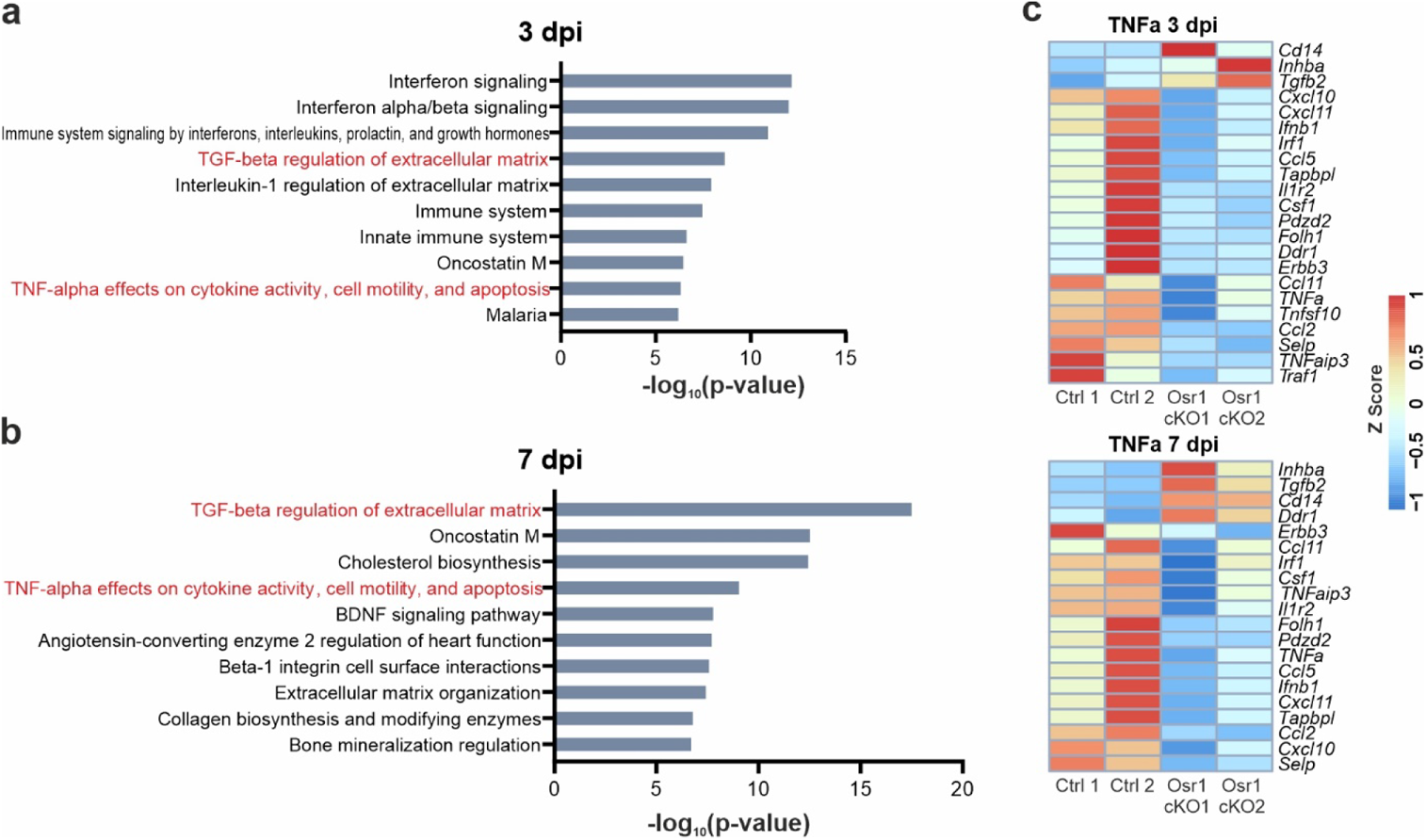
TNFα pathway analysis in transcriptome data of Osr1cKO FAPs. **a, b** GO term analysis of DE genes in Osr1cKO FAPs relative to control FAPs at 3 dpi (top) and 7 dpi (bottom). **c** Heat maps showing downregulation of TNFα pathway genes belonging to the highlighted terms in Osr1cKO FAPs.

**Figure S7.**
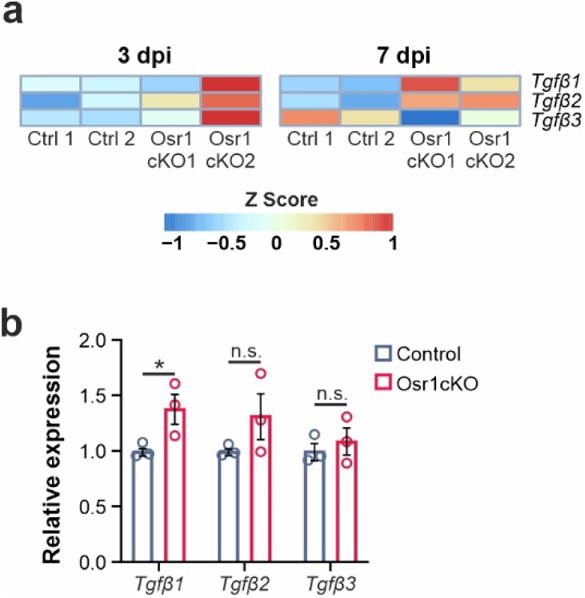
Upregulation of *Tgfb* genes in Osr1cKO FAPs. **a** Heat maps showing *Tgfb1,2* and *3* gene expression in control or Osr1cKO FAPs at 3 and 7 dpi. **b** RT-qPCR analysis of *Tgfb1,2* and *3* gene expression in *in vitro* recombined Osr1cKO FAPs (n=3). Data are mean ± SEM; P-value calculated by Mann-Whitney test; * p < 0.05. N-numbers indicate biological replicates (mice per genotype).

**Figure S8.**
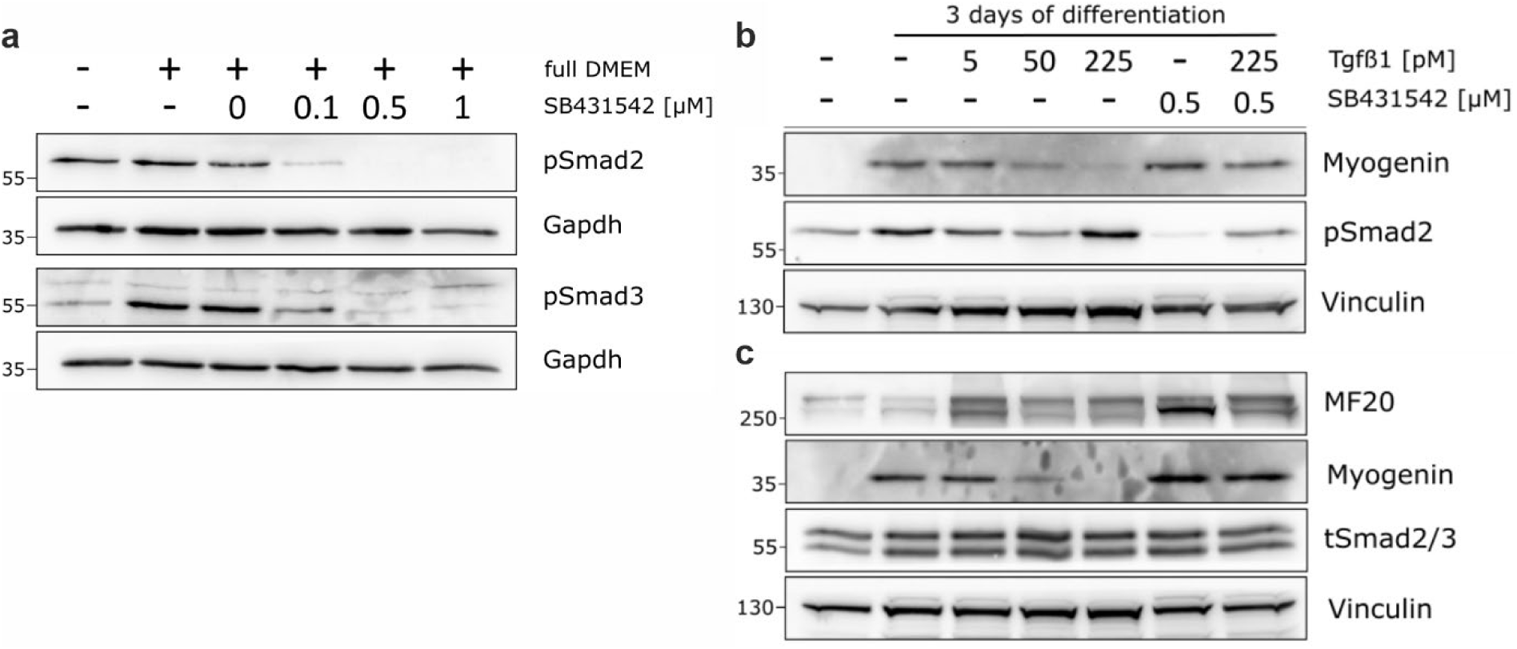
Efficacy of TGFβ pathway inhibition and upregulation of Myogenin by SB431542 in C2C12 cells. **a** Western blot analysis of TGFβ pathway activity in C2C12 cells cultured in DMEM and treated with increasing concentrations of SB4315242 assessed by detection of phosphor-Smad2 phosphor-Smad3. **b, c** Western blot analysis of phospho-Smad2, Myogenin and myosin heavy chain (MF20) expression in C2C12 cells treated with recombinant TGFβ1 and SB4315242.

**Figure S9.**
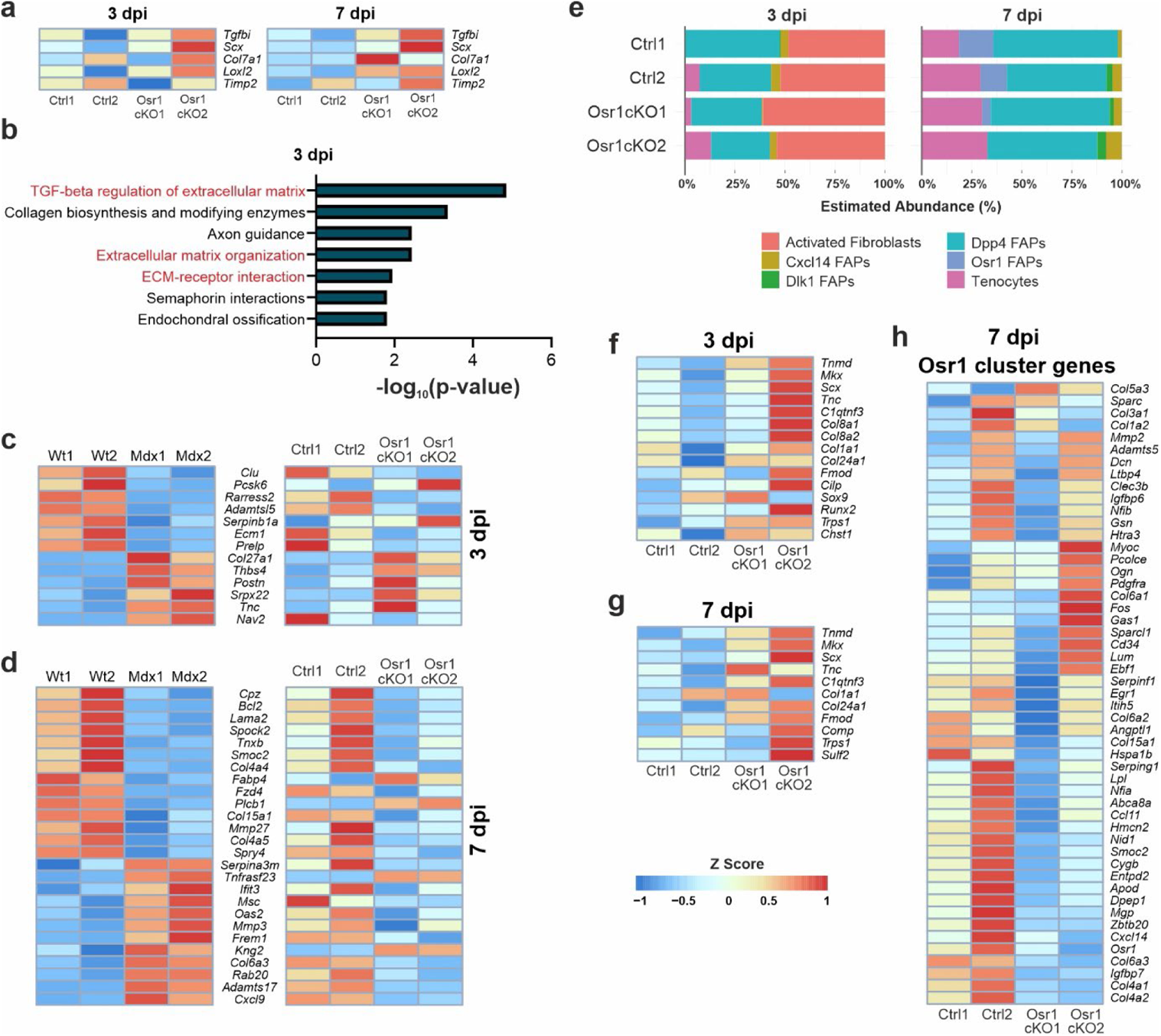
Transcriptional fibrogenic shift of Osr1cKO FAPs. **a** Heat maps showing TGFβ target gene expression in control or Osr1cKO FAPs at 3 and 7 dpi. **b** Bio planet 2019 pathway analysis of genes upregulated in 7 dpi Osr1cKO FAPs relative to controls. **c, d** Heat maps showing in part overlapping ECM gene deregulation between mdx FAPs and 3 dpi (b) and 7 dpi (c) Osr1cKO FAPs. **e** Deconvolution analysis of control and Osr1cKO FAP bulk transcriptome data on single cell sequencing data from Oprescu et al. **f, g** Heat maps showing upregulation of tendon- and cartilage-associated genes in Osr1cKO FAPs relative to controls. **h** Heat map depiction of signature genes characterizing the “*Osr1*” cluster in Oprescu et al.

**Figure S10.**
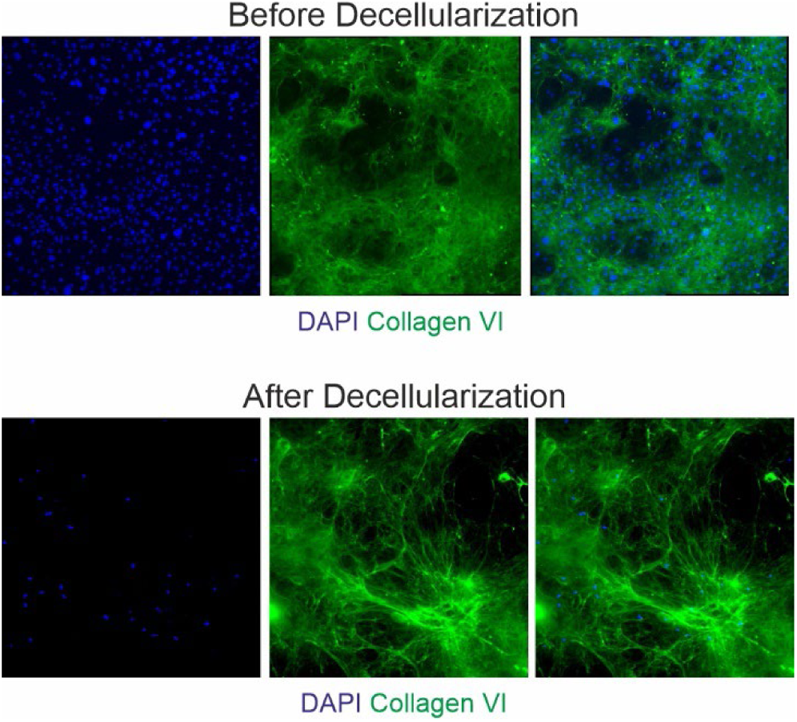
Efficiency of in vitro ECM deposition and the decellularization.

### Supplementary Tables

**Table S1.**
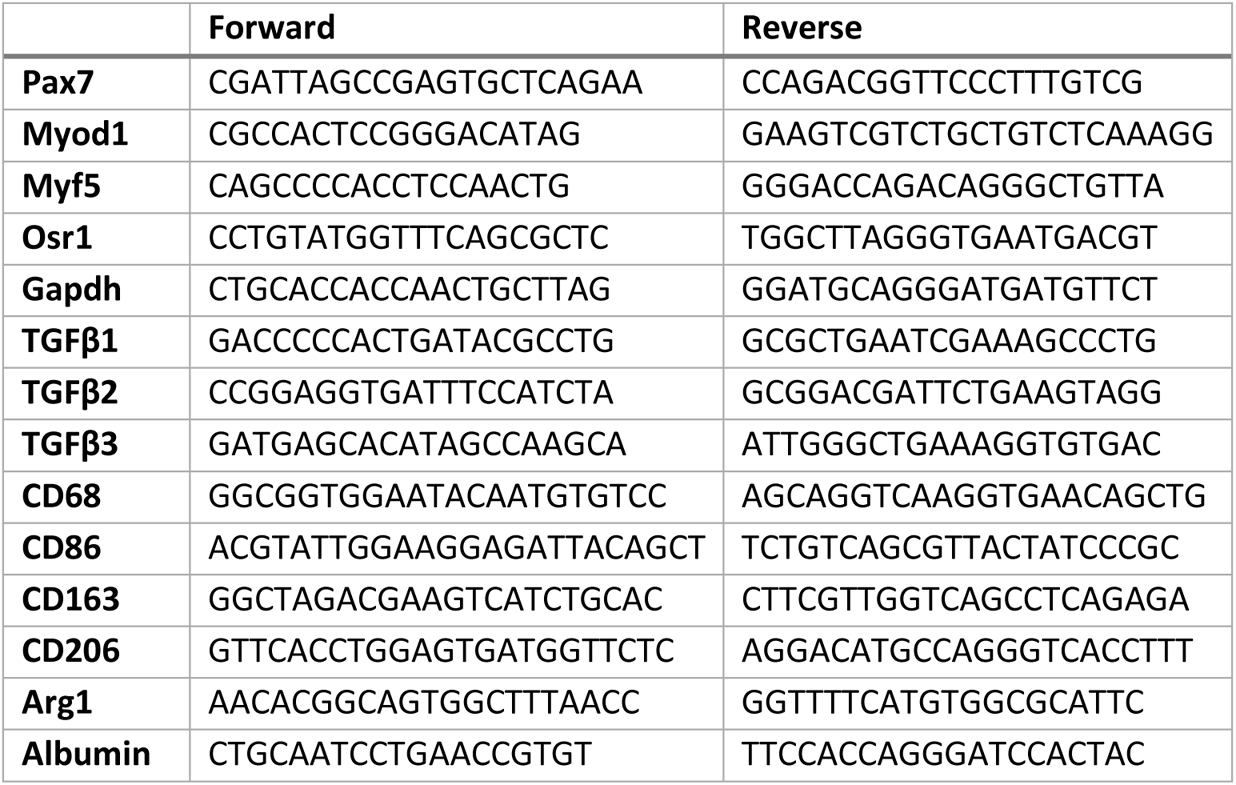
Primers for RT-qPCR.

**Table S2.** Primary antibodies.

| Antibody | Clone | Conjugate | Concentration / Dilution | Source |
| --- | --- | --- | --- | --- |
| **Mouse anti-MHC3 (eMHC)** | Monoclonal | Unconjugated | 1:50 | DSHB |
| **Mouse anti-MF20 (MYH1)** | Monoclonal | Unconjugated | 1:100 | DSHB |
| **Mouse anti-Pax7 (supernatant)** | Monoclonal | Unconjugated | 1:20 | DSHB |
| **Guinea pig anti-Pax7** | Polyclonal | Unconjugated | 1:100 | C. Birchmeier |
| **Mouse anti-Ki67** | Monoclonal | B56 | 1:100 | BD Biosciences |
| **Rabbit anti-Ki67** | Polyclonal | Unconjugated | 1:1000 | Abcam |
| **Rabbit anti-laminin** | Polyclonal | Unconjugated | 5 µg ml<sup>-1</sup> | Sigma-Aldrich |
| **Goat anti-Collagen VI** | Polyclonal | Unconjugated | 1:200 | Southern Biotechnology Associates |
| **Mouse anti-MyoD** | 5.8A | Unconjugated | 1:100 | BD Biosciences |
| **Mouse anti-fibronectin** | Polyclonal | Unconjugated | 1:500 | Merck |
| **Phalloidin** |  | 591/608 | 1:250 | Thermo Fischer Scientific |
| **Rabbit anti-tSmad2/3** | D7G7 | Unconjugated | 1:1000 | Cell Signaling |
| **Rabbit anti-pSmad2** | Ser465/467 | Unconjugated | 1:1000 | Cell Signaling |
| **Rabbit anti-pSmad3** | Ser423/425 | Unconjugated | 1:1000 | Cell Signaling |
| **Rabbit anti-GAPDH** | 14C10 | Unconjugated | 1:2000 | Cell Signaling |
| **Mouse anti-myogenin** | sc-12732 | Unconjugated | 1:100 | Santa Cruz |
| **Mouse anti-vinculin** | V9131 | Unconjugated | 1:4000 | Merck |

**Table S3.** Secondary antibodies.

| **Antibody** | **Conjugate(s)** | **Source** |
| --- | --- | --- |
| Donkey anti-mouse | Alexa Fluor 488, 568 and 680 | Invitrogen |
| Donkey anti-rabbit | Alexa Fluor 488, 568 and 681 | Invitrogen |
| Donkey anti-goat | Alexa Fluor 488, 568 and 682 | Invitrogen |
| Goat anti-guinea pig | Alexa Fluor 682 | Invitrogen |

**Table S4.** FACS antibodies.

| **Antibody** | **Clone** | **Conjugate** | **Concentration/dilution** | **Source** |
| --- | --- | --- | --- | --- |
| **Anti mouse CD31** | 390 | APC conjugated | 3 µg ml<sup>-1</sup> | Invitrogen |
| **Anti mouse CD45** | 30-F11 | APC conjugated | 6 µg ml<sup>-1</sup> | Invitrogen |
| **Anti mouse TER119** | TER-119 | APC conjugated | 6 µg ml<sup>-1</sup> | Invitrogen |
| **Anti rat α7-integrin** | R2F2 | PE conjugated | 1.5 µg ml<sup>-1</sup> | Ablab |
| **Anti mouse Ly-6A/E (Sca-1)** | D7 | APC-Cy7 conjugated | 1.5 µg ml<sup>-1</sup> | Biolegend |
| **CD45** | 30-F11 | AF488 | 0.25 µg ml<sup>-1</sup> | BioLegend, San Diego, CA, USA |
| **CD3e** | 145-2C11 | PE-Cy7 | 0.4 µg ml<sup>-1</sup> | BioLegend, San Diego, CA, USA |
| **CD19** | 6D5 | PE-Cy7 | 0.2 µg ml<sup>-1</sup> | BioLegend, San Diego, CA, USA |
| **CD335** | 29A1.4 | PE-Cy7 | 0.6 µg ml<sup>-1</sup> | BioLegend, San Diego, CA, USA |
| **CD11c** | N418 | PerCP-Cy5.5 | 0.4 µg ml<sup>-1</sup> | BioLegend, San Diego, CA, USA |
| **CD11b/Mac-1** | M1/70 | BV510 | 0.4 µg ml<sup>-1</sup> | BioLegend, San Diego, CA, USA |
| **Ly6-G** | 1A8 | APC-Fire750 | 0.6 µg ml<sup>-1</sup> | BioLegend, San Diego, CA, USA |
| **MHC class II** | M5/114.15.2 | BV785 | 0.05 µg ml<sup>-1</sup> | BioLegend, San Diego, CA, USA |
| **CD80** | 16-10A1 | BV650 | 0.2 µg ml<sup>-1</sup> | BioLegend, San Diego, CA, USA |
| **CD86** | GL-1 | BV421 | 0.2 µg ml<sup>-1</sup> | BioLegend, San Diego, CA, USA |
| **CD163** | TNKUPJ | PE | 0.1 µg ml<sup>-1</sup> | ThermoFisher, Waltham, MA, USA |
| **VEGF** | VG1 | AF647 | 0.225 µg ml<sup>-1</sup> | Novus Biologicals, Littleton, CO, USA |
| **CD206** | C068C2 | PE-Dazzle594 | 0.2 µg ml<sup>-1</sup> | BioLegend, San Diego, CA, USA |

## References

1. Qazi, T.H., et al. Cell therapy to improve regeneration of skeletal muscle injuries. Journal of Cachexia, Sarcopenia and Muscle 10, 501–516 (2019).

2. Sambasivan, R., et al. Pax7-expressing satellite cells are indispensable for adult skeletal muscle regeneration. Development 138, 3647–3656 (2011).

3. Murphy, M.M., Lawson, J.A., Mathew, S.J., Hutcheson, D.A. & Kardon, G. Satellite cells, connective tissue fibroblasts and their interactions are crucial for muscle regeneration. Development 138, 3625–3637 (2011).

4. Dumont, N.A., Wang, Y.X. & Rudnicki, M.A. Intrinsic and extrinsic mechanisms regulating satellite cell function. Development 142, 1572 (2015).

5. Motohashi, N. & Asakura, A. Muscle satellite cell heterogeneity and self-renewal. Front Cell Dev Biol 2, 1 (2014).

6. Wosczyna, M.N. & Rando, T.A. A Muscle Stem Cell Support Group: Coordinated Cellular Responses in Muscle Regeneration. Dev Cell 46, 135–143 (2018).

7. Saclier, M., Cuvellier, S., Magnan, M., Mounier, R. & Chazaud, B. Monocyte/macrophage interactions with myogenic precursor cells during skeletal muscle regeneration. FEBS J 280, 4118–4130 (2013).

8. Arnold, L., et al. Inflammatory monocytes recruited after skeletal muscle injury switch into antiinflammatory macrophages to support myogenesis. J Exp Med 204, 1057–1069 (2007).

9. Chazaud, B. Inflammation and Skeletal Muscle Regeneration: Leave It to the Macrophages! Trends in Immunology 41, 481–492 (2020).

10. Heredia, J.E., et al. Type 2 innate signals stimulate fibro/adipogenic progenitors to facilitate muscle regeneration. Cell 153, 376–388 (2013).

11. Joe, A.W., et al. Muscle injury activates resident fibro/adipogenic progenitors that facilitate myogenesis. Nat Cell Biol 12, 153–163 (2010).

12. Uezumi, A., Fukada, S.-I., Yamamoto, N., Takeda, S.I. & Tsuchida, K. Mesenchymal progenitors distinct from satellite cells contribute to ectopic fat cell formation in skeletal muscle. Nature Cell Biology 12, 143–152 (2010).

13. Uezumi, A., Ikemoto-Uezumi, M. & Tsuchida, K. Roles of nonmyogenic mesenchymal progenitors in pathogenesis and regeneration of skeletal muscle. Front Physiol 5, 68 (2014).

14. Uezumi, A., et al. Fibrosis and adipogenesis originate from a common mesenchymal progenitor in skeletal muscle. J Cell Sci 124, 3654–3664 (2011).

15. Fiore, D., et al. Pharmacological blockage of fibro/adipogenic progenitor expansion and suppression of regenerative fibrogenesis is associated with impaired skeletal muscle regeneration. Stem Cell Research 17, 161–169 (2016).

16. Wosczyna, M.N., et al. Mesenchymal Stromal Cells Are Required for Regeneration and Homeostatic Maintenance of Skeletal Muscle. Cell Rep 27, 2029–2035 e2025 (2019).

17. Hogarth, M.W., et al. Fibroadipogenic progenitors are responsible for muscle loss in limb girdle muscular dystrophy 2B. Nature Communications 10 (2019).

18. Madaro, L., et al. Denervation-activated STAT3-IL-6 signalling in fibro-adipogenic progenitors promotes myofibres atrophy and fibrosis. Nat Cell Biol 20, 917–927 (2018).

19. Kopinke, D., Roberson, E.C. & Reiter, J.F. Ciliary Hedgehog Signaling Restricts Injury-Induced Adipogenesis. Cell 170, 340–351.e312 (2017).

20. Agley, C.C., Rowlerson, A.M., Velloso, C.P., Lazarus, N.R. & Harridge, S.D.R. Human skeletal muscle fibroblasts, but not myogenic cells, readily undergo adipogenic differentiation. Journal of Cell Science 126, 5610–5625 (2013).

21. Contreras, O., Rebolledo, D.L., Oyarzún, J.E., Olguín, H.C. & Brandan, E. Connective tissue cells expressing fibro/adipogenic progenitor markers increase under chronic damage: relevance in fibroblast-myofibroblast differentiation and skeletal muscle fibrosis. Cell and Tissue Research 364, 647–660 (2016).

22. Lemos, D.R., et al. Nilotinib reduces muscle fibrosis in chronic muscle injury by promoting TNF-mediated apoptosis of fibro/adipogenic progenitors. Nat Med 21, 786–794 (2015).

23. Gonzalez, D., et al. ALS skeletal muscle shows enhanced TGF-β signaling, fibrosis and induction of fibro/adipogenic progenitor markers. PLOS ONE 12, e0177649 (2017).

24. Vallecillo-Garcia, P., et al. Odd skipped-related 1 identifies a population of embryonic fibro-adipogenic progenitors regulating myogenesis during limb development. Nat Commun 8, 1218 (2017).

25. Stumm, J., et al. Odd skipped-related 1 (Osr1) identifies muscle-interstitial fibro-adipogenic progenitors (FAPs) activated by acute injury. Stem Cell Res 32, 8–16 (2018).

26. Oprescu, S.N., Yue, F., Qiu, J., Brito, L.F. & Kuang, S. Temporal Dynamics and Heterogeneity of Cell Populations during Skeletal Muscle Regeneration. iScience 23, 100993 (2020).

27. Yang, S. & Plotnikov, S.V. Mechanosensitive Regulation of Fibrosis. Cells 10, 994 (2021).

28. Brashear, S.E., Wohlgemuth, R.P., Gonzalez, G. & Smith, L.R. Passive stiffness of fibrotic skeletal muscle in mdx mice relates to collagen architecture. The Journal of Physiology 599, 943–962 (2021).

29. Kiriaev, L., et al. Lifespan analysis of dystrophic mdx fast-twitch muscle morphology and its impact on contractile function. (Cold Spring Harbor Laboratory, 2021).

30. Smith, L.R. & Barton, E.R. Regulation of fibrosis in muscular dystrophy. Matrix Biology 68-69, 602–615 (2018).

31. Contreras, O., Rossi, F.M. & Brandan, E. Adherent muscle connective tissue fibroblasts are phenotypically and biochemically equivalent to stromal fibro/adipogenic progenitors. Matrix Biology Plus 2 (2019).

32. Helmbacher, F. & Stricker, S. Tissue cross talks governing limb muscle development and regeneration. Seminars in Cell & Developmental Biology 104, 14–30 (2020).

33. Rodgers, J.T., et al. mTORC1 controls the adaptive transition of quiescent stem cells from G0 to GAlert. Nature 510, 393–396 (2014).

34. Massague, J., Cheifetz, S., Endo, T. & Nadal-Ginard, B. Type beta transforming growth factor is an inhibitor of myogenic differentiation. Proceedings of the National Academy of Sciences 83, 8206–8210 (1986).

35. Olson, E.N., Sternberg, E., Hu, J.S., Spizz, G. & Wilcox, C. Regulation of myogenic differentiation by type beta transforming growth factor. The Journal of Cell Biology 103, 1799–1805 (1986).

36. Girardi, F., et al. TGFbeta signaling curbs cell fusion and muscle regeneration. Nat Commun 12, 750 (2021).

37. MacDonald, E.M. & Cohn, R.D. TGFβ signaling: its role in fibrosis formation and myopathies. Current opinion in rheumatology 24, 628–634 (2012).

38. Bensalah, M., et al. A negative feedback loop between fibroadipogenic progenitors and muscle fibres involving endothelin promotes human muscle fibrosis. Journal of Cachexia, Sarcopenia and Muscle n/a (2022).

39. Mendias, C.L., et al. Transforming growth factor-beta induces skeletal muscle atrophy and fibrosis through the induction of atrogin-1 and scleraxis. Muscle & nerve 45, 55–59 (2012).

40. Vindevoghel, L., et al. Smad-dependent transcriptional activation of human type VII collagen gene (COL7A1) promoter by transforming growth factor-β. Journal of Biological Chemistry 273, 13053–13057 (1998).

41. Sethi, A., Mao, W., Wordinger, R.J. & Clark, A.F. Transforming growth factor-beta induces extracellular matrix protein cross-linking lysyl oxidase (LOX) genes in human trabecular meshwork cells. Invest Ophthalmol Vis Sci 52, 5240–5250 (2011).

42. Malecova, B., et al. Dynamics of cellular states of fibro-adipogenic progenitors during myogenesis and muscular dystrophy. Nat Commun 9, 3670 (2018).

43. Wang, X., Park, J., Susztak, K., Zhang, N.R. & Li, M. Bulk tissue cell type deconvolution with multi-subject single-cell expression reference. Nature communications 10, 1–9 (2019).

44. Lukjanenko, L., et al. Aging Disrupts Muscle Stem Cell Function by Impairing Matricellular WISP1 Secretion from Fibro-Adipogenic Progenitors. Cell Stem Cell (2019).

45. Bosnakovski, D., et al. Muscle pathology from stochastic low level DUX4 expression in an FSHD mouse model. Nature Communications 8 (2017).

46. Lapidos, K.A., Kakkar, R. & McNally, E.M. The Dystrophin Glycoprotein Complex. Circulation Research 94, 1023–1031 (2004).

47. Sandonà, M., et al. HDAC inhibitors tune miRNAs in extracellular vesicles of dystrophic muscle-resident mesenchymal cells. EMBO reports 21 (2020).

48. Juban, G., et al. AMPK Activation Regulates LTBP4-Dependent TGF-beta1 Secretion by Pro-inflammatory Macrophages and Controls Fibrosis in Duchenne Muscular Dystrophy. Cell Rep 25, 2163–2176 e2166 (2018).

49. Giordani, L., et al. High-Dimensional Single-Cell Cartography Reveals Novel Skeletal Muscle-Resident Cell Populations. Molecular Cell 74, 609–621.e606 (2019).

50. Scott, R.W., Arostegui, M., Schweitzer, R., Rossi, F.M.V. & Underhill, T.M. Hic1 Defines Quiescent Mesenchymal Progenitor Subpopulations with Distinct Functions and Fates in Skeletal Muscle Regeneration. Cell Stem Cell 25, 797–813 e799 (2019).

51. De Micheli, A.J., et al. Single-Cell Analysis of the Muscle Stem Cell Hierarchy Identifies Heterotypic Communication Signals Involved in Skeletal Muscle Regeneration. Cell Reports 30, 3583–3595.e3585 (2020).

52. De Micheli, A.J., Spector, J.A., Elemento, O. & Cosgrove, B.D. A reference single-cell transcriptomic atlas of human skeletal muscle tissue reveals bifurcated muscle stem cell populations. Skeletal Muscle 10, 19 (2020).

53. Dey, D., et al. Two tissue-resident progenitor lineages drive distinct phenotypes of heterotopic ossification. Science Translational Medicine 8, 366ra163-366ra163 (2016).

54. Lees-Shepard, J.B., et al. Activin-dependent signaling in fibro/adipogenic progenitors causes fibrodysplasia ossificans progressiva. Nature Communications 9 (2018).

55. Forcina, L., Cosentino, M. & Musarò, A. Mechanisms Regulating Muscle Regeneration: Insights into the Interrelated and Time-Dependent Phases of Tissue Healing. Cells 9 (2020).

56. Saclier, M., Cuvellier, S., Magnan, M., Mounier, R. & Chazaud, B. Monocyte/macrophage interactions with myogenic precursor cells during skeletal muscle regeneration. FEBS Journal 280, 4118–4130 (2013).

57. Sass, F., et al. Immunology Guides Skeletal Muscle Regeneration. International Journal of Molecular Sciences 19, 835 (2018).

58. Roca, H., et al. CCL2 and Interleukin-6 Promote Survival of Human CD11b+ Peripheral Blood Mononuclear Cells and Induce M2-type Macrophage Polarization. Journal of Biological Chemistry 284, 34342–34354 (2009).

59. Sierra-Filardi, E., et al. CCL2 Shapes Macrophage Polarization by GM-CSF and M-CSF: Identification of CCL2/CCR2-Dependent Gene Expression Profile. The Journal of Immunology 192, 3858–3867 (2014).

60. Farmaki, E., Kaza, V., Chatzistamou, I. & Kiaris, H. CCL8 Promotes Postpartum Breast Cancer by Recruiting M2 Macrophages. iScience 23, 101217 (2020).

61. Tripathi, C., et al. Macrophages are recruited to hypoxic tumor areas and acquire a Pro-Angiogenic M2-Polarized phenotype via hypoxic cancer cell derived cytokines Oncostatin M and Eotaxin. Oncotarget 5, 5350–5368 (2014).

62. Shirakawa, T., et al. Tumor necrosis factor alpha regulates myogenesis to inhibit differentiation and promote proliferation in satellite cells. Biochem Biophys Res Commun 580, 35–40 (2021).

63. Urciuolo, A., et al. Collagen VI regulates satellite cell self-renewal and muscle regeneration. Nat Commun 4, 1964 (2013).

64. Andenæs, K., et al. The extracellular matrix proteoglycan fibromodulin is upregulated in clinical and experimental heart failure and affects cardiac remodeling. PLOS ONE 13, e0201422 (2018).

65. Pourhanifeh, M.H., et al. The role of fibromodulin in cancer pathogenesis: implications for diagnosis and therapy. Cancer Cell International 19 (2019).

66. Schumacher, A., et al. Angptl4 is upregulated under inflammatory conditions in the bone marrow of mice, expands myeloid progenitors, and accelerates reconstitution of platelets after myelosuppressive therapy. Journal of Hematology & Oncology 8 (2015).

67. Kadomatsu, T., Endo, M., Miyata, K. & Oike, Y. Diverse roles of ANGPTL2 in physiology and pathophysiology. Trends in Endocrinology & Metabolism 25, 245–254 (2014).

68. Zhao, J., et al. Age-dependent increase in angiopoietin-like protein 2 accelerates skeletal muscle loss in mice. Journal of Biological Chemistry 293, 1596–1609 (2018).

69. Descamps, S., et al. Inhibition of myoblast differentiation by Sfrp1 and Sfrp2. Cell and Tissue Research 332, 299–306 (2008).

70. Sohn, J., Lu, A., Tang, Y., Wang, B. & Huard, J. Activation of non-myogenic mesenchymal stem cells during the disease progression in dystrophic dystrophin/utrophin knockout mice. Human Molecular Genetics (2015).

71. Csapo, R., Gumpenberger, M. & Wessner, B. Skeletal Muscle Extracellular Matrix – What Do We Know About Its Composition, Regulation, and Physiological Roles? A Narrative Review. Frontiers in Physiology 11 (2020).

72. Silver, J.S., et al. Injury-mediated stiffening persistently activates muscle stem cells through YAP and TAZ mechanotransduction. Science Advances 7, eabe4501 (2021).

73. Trensz, F., et al. Increased microenvironment stiffness in damaged myofibers promotes myogenic progenitor cell proliferation. Skeletal Muscle 5 (2015).

74. Cosgrove, B.D., Sacco, A., Gilbert, P.M. & Blau, H.M. A home away from home: Challenges and opportunities in engineering in vitro muscle satellite cell niches. Differentiation 78, 185–194 (2009).

75. Barraza-Flores, P., Bates, C.R., Oliveira-Santos, A. & Burkin, D.J. Laminin and Integrin in LAMA2-Related Congenital Muscular Dystrophy: From Disease to Therapeutics. Front Mol Neurosci 13, 1–1 (2020).

76. Ross, J., et al. Defects in Glycosylation Impair Satellite Stem Cell Function and Niche Composition in the Muscles of the Dystrophic LargemydMouse. STEM CELLS 30, 2330–2341 (2012).

77. Lieber, R.L. & Ward, S.R. Cellular Mechanisms of Tissue Fibrosis. 4. Structural and functional consequences of skeletal muscle fibrosis. American Journal of Physiology-Cell Physiology 305, C241–C252 (2013).

78. Lacraz, G., et al. Increased Stiffness in Aged Skeletal Muscle Impairs Muscle Progenitor Cell Proliferative Activity. PLOS ONE 10, e0136217 (2015).

79. Stearns-Reider, K.M., et al. Aging of the skeletal muscle extracellular matrix drives a stem cell fibrogenic conversion. Aging Cell 16, 518–528 (2017).

80. Hogarth, M.W., Uapinyoying, P., Mázala, D.A.G. & Jaiswal, J.K. Pathogenic role and therapeutic potential of fibro-adipogenic progenitors in muscle disease. Trends in Molecular Medicine (2021).

81. Accorsi, A., Cramer, M.L. & Girgenrath, M. Fibrogenesis in LAMA2-Related Muscular Dystrophy Is a Central Tenet of Disease Etiology. Front Mol Neurosci 13 (2020).

82. Ismaeel, A., et al. Role of Transforming Growth Factor-β in Skeletal Muscle Fibrosis: A Review. International Journal of Molecular Sciences 20, 2446 (2019).

83. Nitahara-Kasahara, Y., et al. Dystrophic mdx mice develop severe cardiac and respiratory dysfunction following genetic ablation of the anti-inflammatory cytokine IL-10. Human Molecular Genetics 23, 3990–4000 (2014).

84. Vidal, B., et al. Fibrinogen drives dystrophic muscle fibrosis via a TGFβ/alternative macrophage activation pathway. Genes & Development 22, 1747–1752 (2008).

85. Pessina, P., et al. Novel and optimized strategies for inducing fibrosis in vivo: focus on Duchenne Muscular Dystrophy. Skeletal Muscle 4, 7 (2014).

86. Biressi, S., Miyabara, E.H., Gopinath, S.D., M. Carlig, P.M. & Rando, T.A. A Wnt-TGFβ2 axis induces a fibrogenic program in muscle stem cells from dystrophic mice. Science Translational Medicine 6, 267ra176-267ra261 (2014).

87. Carlson, M.E., Hsu, M. & Conboy, I.M. Imbalance between pSmad3 and Notch induces CDK inhibitors in old muscle stem cells. Nature 454, 528–532 (2008).

88. Hayashi, S. & McMahon, A.P. Efficient Recombination in Diverse Tissues by a Tamoxifen-Inducible Form of Cre: A Tool for Temporally Regulated Gene Activation/Inactivation in the Mouse. Developmental Biology 244, 305–318 (2002).

89. Rodríguez, C.I., et al. High-efficiency deleter mice show that FLPe is an alternative to Cre-loxP. Nat Genet 25, 139–140 (2000).

90. Bucher, C.H., et al. Experience in the Adaptive Immunity Impacts Bone Homeostasis, Remodeling, and Healing. Frontiers in Immunology 10 (2019).

91. Kuleshov, M.V., et al. Enrichr: a comprehensive gene set enrichment analysis web server 2016 update. Nucleic acids research 44, W90–W97 (2016).

92. Reimand, J., et al. Pathway enrichment analysis and visualization of omics data using g: Profiler, GSEA, Cytoscape and EnrichmentMap. Nature protocols 14, 482–517 (2019).

